# Context-dependent siderophore exploitability shapes microbial community structure

**DOI:** 10.64898/2026.05.27.728356

**Authors:** Aileen Krüger, Seung-Hyun Paik, Michelle Bund, Matthias Pesch, Stephan Krueger, Benita Lückel, Athanasios Papadopoulos, Johanna Wiechert, Tanvir Hassan, Ulrike Weber, Romain Avellan, Sander Smits, Michael Bott, Ákos T. Kovács, Philipp Westhoff, Anna Matuszyńska, Dietrich Kohlheyer, Thomas Drepper, Julia Frunzke

## Abstract

Siderophores are classically viewed as shared iron-scavenging public goods, yet their ecological roles in multispecies communities remain poorly defined. Here, we establish a synthetic microbial community to dissect how different siderophores, their uptake compatibility and spatial structure shape iron competition. Using *Corynebacterium glutamicum* as a model, we show that this siderophore non-producer accesses diverse xenosiderophores, including enterobactin secreted by *Escherichia coli*. However, exploitation was constrained and co-cultures converged to stable compositions. Dose-response experiments combined with mathematical modelling indicated that the producer retains more effective access to enterobactin than the exploiter. Presence of *Pseudomonas putida* altered this interaction, as it exploited enterobactin while producing pyoverdine, a siderophore inaccessible to the other community members that restricted their iron access. Across different cultivation scales, community dynamics was strongly influenced by spatial organization and initial composition. These findings identify siderophores as context-dependent iron-allocation agents that can promote microbial coexistence or exclusion.

## 2) Introduction

Iron is essential for almost all living organisms because it serves as a cofactor in fundamental cellular processes including respiration, central carbon metabolism, and DNA synthesis [1,2]. Yet, despite its abundance in many environments, biologically accessible iron is often scarce under aerobic conditions, where iron occurs predominantly as poorly soluble ferric iron [3]. To overcome this limitation, many bacteria, fungi, and plants produce siderophores that are small high-affinity metabolites that chelate ferric iron and deliver it to cells through dedicated uptake systems [4]. As the synthesis of siderophores requires a significant metabolic effort, they are only synthesized and secreted in response to iron limitation [5,6]. Today, more than 500 siderophores are known, with a broad variety of structural classifications, such as e.g. catecholate, carboxylate, hydroxamate or mixed-type siderophores [7,8]. The archetype and siderophore with the highest affinity for iron reported to date is enterobactin, which is produced by *Escherichia coli* and *Klebsiella pneumoniae* [9].

Siderophores are not only iron-scavenging molecules but also mediators of microbial interactions [2]. In bacterial communities, siderophore-bound iron can be accessed only by organisms carrying compatible uptake systems. As a result, siderophores may function as shared resources, privatized goods, or competitive agents [2]. Gram-negative bacteria recognize specific iron-loaded siderophores via matching outer membrane receptors, like FepA of *Escherichia coli* for enterobactin [10], or FpvA in *Pseudomonas putida* for pyoverdine uptake [11]. In contrast, Gram-positive bacteria scavenge them by specific membrane-anchored siderophore-binding lipoproteins and ABC transporters [12]. Many microbes encode receptors and transporters for siderophores that they do not themselves synthesize, enabling them to exploit xenosiderophores produced by neighbours [13-16]. For example, *E. coli* synthesizes and utilizes enterobactin, but also has the receptor FhuA for ferrichrome making use of a siderophore produced by many fungal species like *Penicillium* spp. or *Ustilago* spp. [17,18]. Moreover, *Pseudomonas aeruginosa* upregulates the siderophore transport system FemIRA when facing mycobactin produced by *Mycobacterium* spp. [19]. Xenosiderophore uptake and utilization reduces the metabolic cost of iron acquisition, but it can also deprive competing organisms of accessible iron [2,20,21]. Conversely, siderophore producers can restrict iron availability to organisms lacking matching receptors [22,23]. These dynamics make siderophores a powerful determinant of microbial competition and coexistence in iron-limited habitats such as soil, the gut, and further host-associated microbiomes [24]. In these environments, xenosiderophore exploitation is emerging as an important ecological mechanism and a potential target for intervention. For example, the soil bacterium *Pseudomonas fluorescens* pirates desferrioxamines and coelichelin from *Streptomyces ambofaciens* [25], whereas in the nasal microbiome, corynebacterial species consume siderophores produced by *Staphylococcus aureus*, thereby reducing *S. aureus* growth [26]. Likewise, several *Pseudomonas spp*. have been shown to produce pyoverdines with significant antimicrobial properties against several human pathogens due to provoking iron limitation [27]. To date, the molecular basis of siderophore production, regulation, and chemical diversity has been extensively studied, whereas their ecological roles in diverse microbial communities are only beginning to be understood [2,28]. Most work has focused either on individual siderophore systems [29,30], on pairwise producer-cheater interactions [23,31,32], or influence on growth and performance of microbial co-cultures [33-35]. In contrast, natural communities often contain multiple species that differ in siderophore production, uptake compatibility, and spatial organization. Under these conditions, the ecological role of a siderophore may not be fixed: the same molecule could promote growth of one community member while excluding another, depending on receptor compatibility, relative abundance, and the spatial structure of the habitat. A tractable synthetic system consisting of model bacteria with defined traits is therefore needed to dissect how sharing, exploitation, and exclusion emerge together.

Here, we established a synthetic community to examine context-dependent siderophore interactions. We chose *Corynebacterium glutamicum* as a model xenosiderophore exploiter because corynebacteria were recently described as common exploiters in natural microbiomes [26]. *C. glutamicum* is a well-established, genetically tractable model organism that does not produce siderophores [36], but exhibits a broad repertoire of putative xenosiderophore transport proteins (Table S1) [37,38]. We first identified diverse bacterial and fungal siderophores, including enterobactin, that support growth of iron-starved *C. glutamicum*. We then built a tripartite community consisting of *C. glutamicum* as an exploiter, *E. coli* as an enterobactin-producing public-good provider, and *Pseudomonas putida* as both an enterobactin exploiter and a pyoverdine (selfish good) producer. By analysing community development and composition across scales from single cells to structured communities on plates, we show that siderophore-mediated interactions are strongly context dependent. Enterobactin acts as a public good that supports the growth of each community member, whereas pyoverdine acts as a selfish good, i.e. a non-exploitable good that restricts iron access from *E. coli* and *C. glutamicum* even in the presence of exploitable enterobactin. A mathematical model parameterized with experimental data suggests that exploiters can utilize the siderophore less efficiently than producers, e.g. due to a substantially lower receptor affinity, constraining exploitation despite access to enterobactin. Spatial structure further modulated these interactions by promoting niche formation for exploiters while also facilitating enterobactin privatization by the producer. Together, these findings highlight the context dependency of siderophore-mediated interactions and underscore the central role of spatial organization in shaping microbial community dynamics.

## 3) Results

### 3.1 Corynebacterium glutamicum exploits chemically diverse xenosiderophores under iron limitation

Given the predicted capacity of the non-siderophore-producing *C. glutamicum* to exploit xenosiderophores (Table S1), we first determined which siderophores support its growth under iron limitation. To this end, we systematically assessed the growth of iron-starved *C. glutamicum* in proximity to microorganisms producing chemically distinct siderophores, including catecholates, hydroxamates and mixed-type siderophores (Figure 1A). As expected, *C. glutamicum* failed to grow on solid iron-restricted media in monoculture (Figure S1). However, robust growth was observed when growing next to enterobactin producing *E. coli* and *Serratia odorifera*, bacillibactin secreting *Bacillus subtilis* and *Bacillus amyloliquefaciens*, as well as ferrichrome producing fungus *Ustilago maydis* (Figure 1A). In contrast, no growth was observed next to *Pseudomonas putida* and *Pseudomonas taiwanensis*, both secreting pyoverdine variants. This xenosiderophore utilization pattern is further supported when analyzing growth in liquid cultures supplemented with 50% lysogeny broth (LB)- based spent medium of selected siderophore producing organisms grown under iron depletion (Figure 1B).

**Figure 1:**
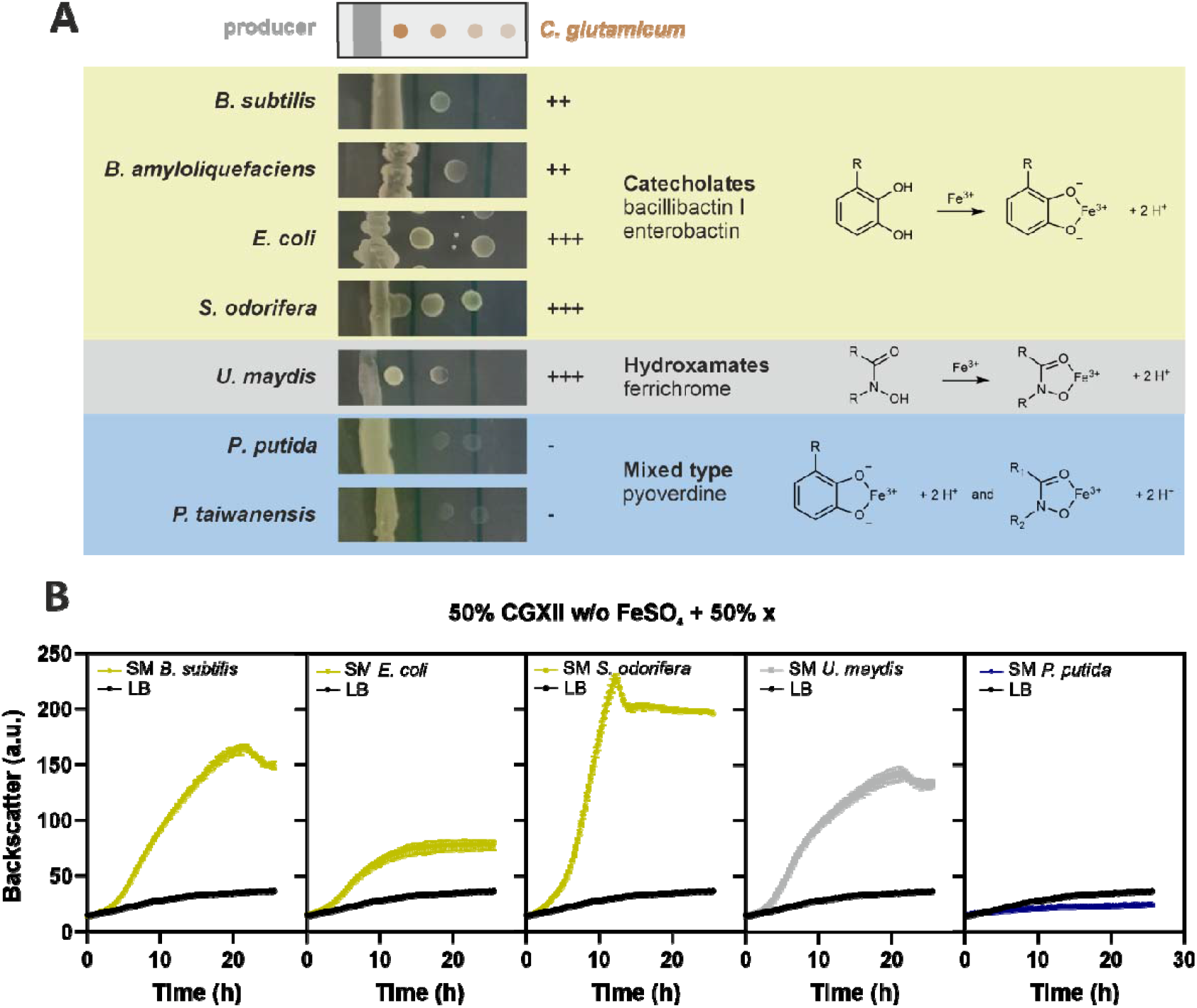
C.*glutamicum* is able to exploit several chemically diverse xenosiderophores. **(A)** The potential of seven known siderophore producers to promote growth of neighboring iron-starved *C. glutamicum* on agar-solidified LB medium containing 250 µM DIP (2,2’-dipyridyl) was investigated. *C. glutamicum* cell suspensions were spotted at different distances to the pre-grown lane of siderophore producer. Number of + and − signs indicate the growth-promoting effect. Moieties involved in iron chelation that determine siderophore classification are shown on the right (adapted from [39]). One representative replicate is shown. **(B)** Liquid cultivation of *C. glutamicum* in 50% 2-fold CGXII minimal media without added iron, and 50% spent medium harvested from selected organisms as in (A) grown in LB supplemented with 200 µM DIP. Graph colors represent siderophore types, while LB control is shown in black. *n* = 3 biological replicates.

Together, these results demonstrate that *C. glutamicum* can exploit a diverse set of xenosiderophores, spanning multiple iron-chelation chemistries and originating from both Gram-positive and Gram-negative bacteria, and even from fungi, to overcome iron limitation.

### 3.2 *C.glutamicum* is an exploiter of enterobactin produced by E. coli

Having demonstrated that *C. glutamicum* can exploit a broad range of xenosiderophores, we next focused on its interaction with enterobactin and its producer *E. coli*. Based on the dose-dependent growth stimulation of *C. glutamicum* observed when supplementing iron-limited medium with LB-based *E. coli* spent medium (SM), a 50% SM dilution supplemented with DIP was used for subsequent experiments (Figure S2A-B).

To test whether this growth-promoting effect specifically depends on enterobactin, we generated an enterobactin deficient *E. coli* strain (Δ*entC*), lacking the gene encoding for the isochorismate synthase EntC that catalyzes the conversion of chorismate to isochorismate in the first step of enterobactin synthesis [40]. In comparison to *E. coli* WT SM, SM of the Δ*entC* mutant did not support growth of C. *glutamicum* (Figure 2A). This effect was also observed for CGXII-based SM, i.e. minimal media (Figure S2C). Using a plasmid-based LacI-P_*tac*_-controlled *entC* expression, the growth-promoting effect could be restored (Figure S2D). Notably, supplementation of pure enterobactin also enhanced growth of iron-starved *C. glutamicum* (Figure 2B).

**Figure 2:**
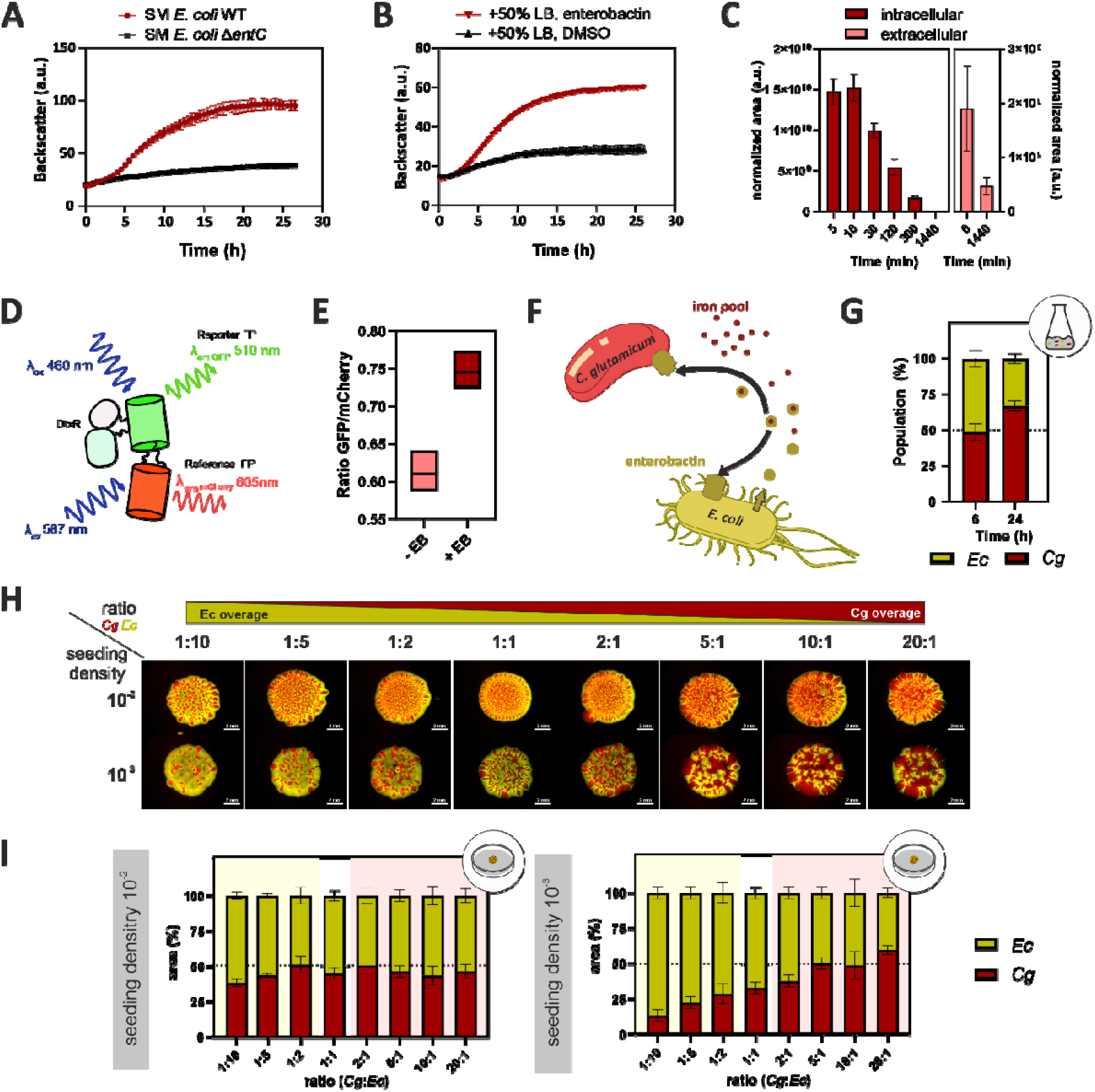
*C. glutamicum* exploits enterobactin produced by *E. coli*. **(A)** Comparison of the growth of *C. glutamicum* in 50% LB-based spent medium of *E. coli* WT (red) with that in spent medium of non-enterobactin-producing *E. coli* Δ*entC* (black). **(B)** *C. glutamicum* growing in 50% LB supplemented with 200 µM DIP and either 10 µM pure enterobactin (red) or 1 µL/mL DMSO as negative control (black). *n* = 3 biological replicates. **(C)** MS-measurement of intra-and extracellular enterobactin content in *C. glutamicum* cultivated with 10 µM enterobactin in the presence of 150 µM DIP. *n* = 4 biological replicates. **(D)** Schematic depiction of ratiometric fluorescent-based modified biosensor “IronSenseR-mCherry” [42]. **(E)** Comparing iron content by utilizing plasmid-based IronSenseR-mCherry in *C. glutamicum* cultivated in CGXII with or without 10 µM enterobactin. **(F)** Schematic depiction of *C. glutamicum* and *E. coli* making use of a non-accessible iron pool via *E. coli-*produced enterobactin. **(G)** Flow cytometry measurements of liquid co-cultured *E. coli-gfp* (yellow) and *C. glutamicum-E2-crimson* (red) under iron-restricted (250 µM DIP) conditions at a ratio of 1:1. Samples taken at 6 h and 24 h after cultivation start. *n* = 3 biological replicates. (H) Representative merged stereomicroscopy images of mixed drop (macrocolony) assay on iron-limited CGXII plates with *C. glutamicum*-*E2-crimson* (red) and *E. coli*-*gfp* (yellow) as indicated in varying ratios and two different seeding densities (10^-2^ and 10^-3^). **(I)** Quantified area of *C. glutamicum* (red) and *E. coli* (yellow) deduced from the microscopy pictures in **(H)**, set in relation to 100%. *n* = 5 biological replicates. *Ec* = *E. coli, Cg* = *C. glutamicum*.

We next asked whether enterobactin is directly taken up by *C. glutamicum* and thereby relieves intracellular iron limitation. Using liquid chromatography–mass spectrometry (LC-MS) measurements intracellular enterobactin was readily detectable immediately after addition, while declined over time, indicating rapid uptake under iron limitation (Figure 2C). However, this fraction may also contain cell-associated enterobactin and does by itself not confirm the uptake of the molecule. Moreover, relative to iron-replete conditions, intracellular citrate levels showed a reduced log_2_-fold change under enterobactin supplementation compared with the no-iron control, suggesting that enterobactin availability alters citrate-associated iron acquisition in *C. glutamicum* [41] (Figure S3). This is also reflected in further TCA cycle intermediates like aconitate and isocitric acid. These results further indicate relieved intracellular iron limitation in the presence of the siderophore. Meanwhile, extracellular enterobactin levels decreased over time, but significant amounts were still detectable after 24 h, showing that enterobactin is not fully degraded by *C. glutamicum* under the tested conditions (Figure 2C).

To further assess enterobactin-mediated intracellular iron availability, the recently described DtxR-based, ratiometric fluorescent iron biosensor (IronSenseR) [42], was modified for *in vivo* measurements by exchanging the Large-stokes shift reference domain (LSSmApple) by a more rapidly maturating fluorescent protein (mCherry) (Figure 2D) and was first validated *in vitro* (Figure S4). The modified IronSenseR variant indicated binding to ferrous iron in low micromolar range as it was observed for the original IronSenseR previously [42] thus indicating that the modification did not affect the biosensor function nor its specificity for ferrous iron (Figure S4). Using this biosensor variant (IronSenseR-mCherry), we observed increased intracellular iron levels upon addition of pure enterobactin, as indicated by an increased GFP/mCherry ratio (Figure 2E). Together, these results demonstrate that enterobactin is directly exploited by *C. glutamicum* and relieves intracellular iron limitation (Figure 2F).

### 3.3 Enterobactin exploitation supports stable pairwise coexistence

As we could demonstrate that *C. glutamicum* is a versatile xenosiderophore exploiter, we next asked how *E. coli*-secreted enterobactin exploitation by *C. glutamicum* unfolds in *in vivo* co-cultivations. To address this, *C. glutamicum* and *E. coli* strains were labelled with constitutively expressed fluorescence reporters E2-Crimson and GFP, respectively, and first analyzed in liquid microtiter cultivations using flow cytometry (Figure 2G). Under iron-restricted conditions and a starting ratio of 1:1, the organisms likewise converged towards approximately equal coexistence, with a slight overrepresentation of *C. glutamicum* after 24 h. Within liquid cultivations, enterobactin availability is equally distributed and sustained cell-to-cell contacts are limited due to shaking. We therefore extended the analysis to spatially structured co-cultures, using macrocolonies in which access to siderophores is likely shaped by cell density, spatial distance, and uptake efficiency. To this end, plate-spot assays were performed under both iron-restricted and iron-replete conditions (Figure S5).

As controls, *E. coli* WT grew in monoculture under both iron-restricted (3.6 µM FeSO_4_ + 100 µM DIP) and iron-replete (36 µM FeSO_4_) conditions. In contrast, neither *C. glutamicum* nor *E. coli* Δ*entC* showed growth under iron limitation (Figure S5A-B). We then co-cultivated *C. glutamicum* with either *E. coli* WT or *E. coli* Δ*entC* on iron-restricted CGXII agar plates at different starting ratios and two seeding densities (10^-2^ and 10^-3^) (Figure 2H) as initial cell densities could influence exploitation and therefore final community composition [43]. As expected, there was no growth when *C. glutamicum* and *E. coli* Δ*entC* were co-cultivated under iron limitation (Figure S5C). In co-culture with *E. coli* WT, prominent growth of *C. glutamicum* was observed across all inoculation ratios, confirming that *C. glutamicum* depends on producer-derived enterobactin also in spatially structured environments. Remarkably, the two species grew in intricate spatial arrangements within the macrocolonies, leading to the formation of distinct red or yellow fluorescent patches irrespective of the ratio and seeding density. Furthermore, quantification of species-specific colony areas revealed that enterobactin exploitation remained markedly constrained: At the higher seeding density (10^-2^), which corresponds to closer spatial proximity, co-cultures converged to an approximately balanced (1:1) outcome across most starting ratios, with only a slight shift toward *E. coli* when it was initially overrepresented (Figure 2I). At lower seeding densities (10^-3^), this balance was less robust when *C. glutamicum* was inoculated at equal or lower abundance, consistent with stronger spatial segregation and reduced access to diffusible enterobactin. Notably, even a 20-fold excess of *C. glutamicum* did not result in dominance over *E. coli*, indicating that the producer still limits the extent of exploitation.

These results suggest that enterobactin is not equally accessible to both partners in macrocolonies like it is the case for the liquid cultivation. Instead, *C. glutamicum* appears to benefit only from the fraction of enterobactin that exceeds recapture by the producer, implying that exploitation depends on the presence of an exploitable surplus. This interpretation is supported by the comparison to *E. coli* WT/*E. coli* Δ*entC* co-cultures, in which both strains share the same uptake system and grew largely in proportion to their initial inoculation ratio, independent of seeding density (Figure S5D). However, when the Δ*entC* mutant is present in excess, a clear difference from *E. coli* WT/*C. glutamicum* macrocolonies could be observed. While the proportion of *C. glutamicum* never exceeded a level of approximately 50%, the excess of the enterobactin-deficient *E. coli* strain retained the initial ratio of the siderophore producer (*E. coli* WT). This observation suggests that the efficiency of enterobactin utilization and/or uptake is significantly lower in *C. glutamicum* than in the *E. coli* Δ*entC* strain.

### 3.4 Single-cell analysis and modelling reveal siderophore-receptor affinity-constrained exploitation in pairwise co-culture

To resolve the dynamics underlying constrained enterobactin exploitation at the single cell level with higher spatiotemporal resolution, we next analysed co-cultivation of fluorescently labelled *E. coli* and *C. glutamicum* as single-layer microcolonies in microfluidic chambers under iron-restricted conditions (Figure 3A, Table S2, Video S1-2). As confirmed previously, *C. glutamicum* grew only in the presence of enterobactin-producing *E. coli* (Figure 3B), but neither in monoculture nor in co-culture with *E. coli* Δ*entC* (Figure 3C). These differences were also reflected in the temporal development of total single-cell area obtained by image-based cell segmentation (Figure 3D).

**Figure 3:**
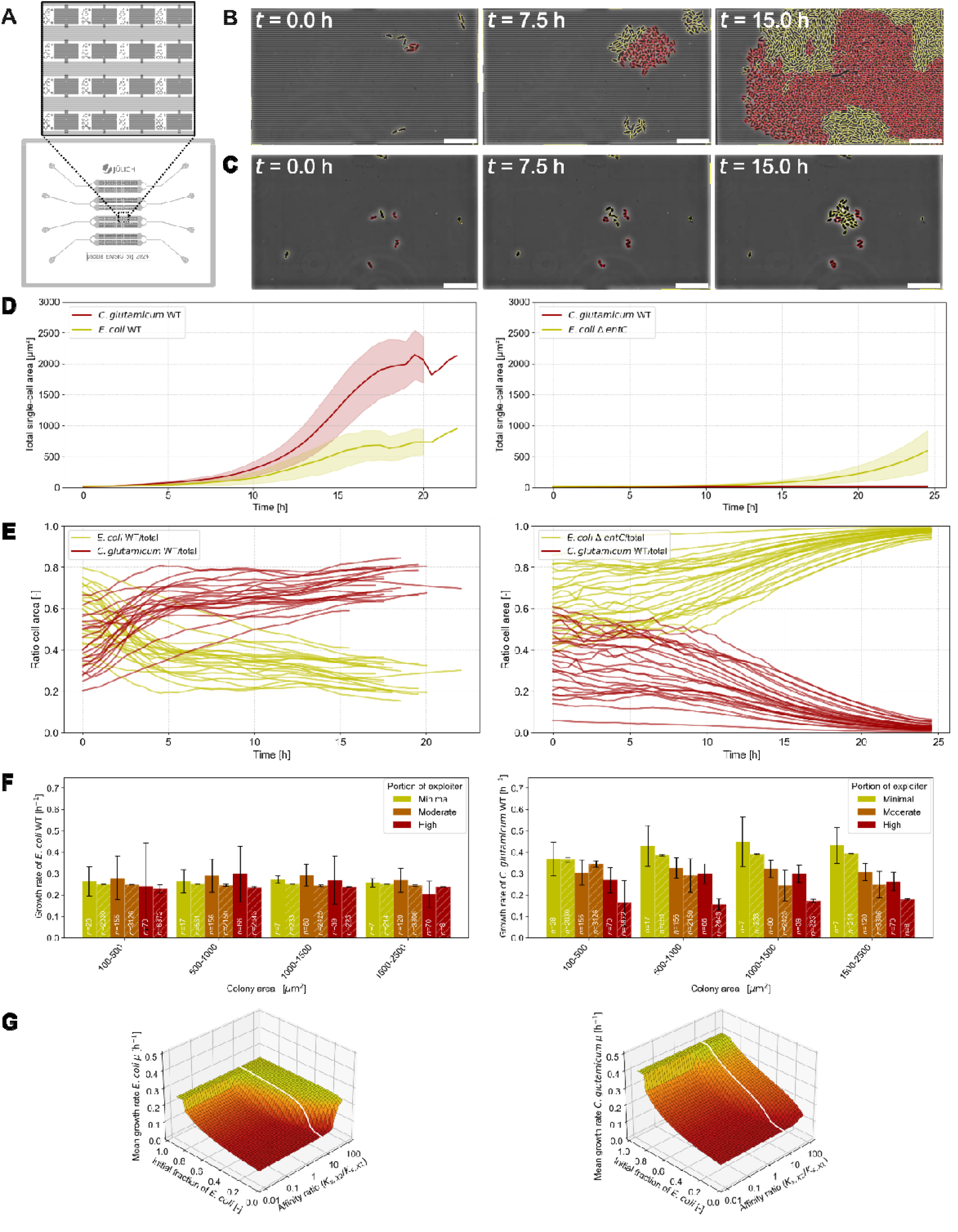
Single-cell analysis and modelling reveal affinity-constrained exploitation in pairwise co-culture of *C. glutamicum* and *E. coli*. **(A)** Schematic layout of the microfluidic device used for co-cultivation. **(B)** Representative segmented frames of the co-culture of *C. glutamicum* WT (red) and *E. coli* WT (yellow) at 0, 7.5 and 15 h in CGXII medium under iron limitation (3.6 µM FeSO_4_) at 30 °C. **(C)** Representative segmented frames of the co-culture of *C. glutamicum* WT (red) and *E. coli* Δ*entC* (yellow) at 0, 7.5 and 15 h under identical conditions. **(D)** Mean total single-cell area (µm^2^) over time. Left, *C. glutamicum* WT (red) and *E. coli* WT (yellow) with *N* = 24; right, *C. glutamicum* WT (red) and *E. coli* Δ*entC* (yellow) with *N* = 27. **(E)** Temporal development of cell area shown separately for each organism. Each line represents an individual chamber. Left, *C. glutamicum* WT (red) and *E. coli* WT (yellow) with *N* = 24; right, *C. glutamicum* WT (red) and *E. coli* Δ*entC* (yellow) with *N* = 27. **(F)** Experimental and model-derived growth-rate dependencies on colony area, fitted with species-specific parameters across colony sizes and exploiter fractions (minimal, 0-0.33; moderate, 0.33-0.66; high, >0.66; color-coded). Dashed lines indicate simulated data, solid lines experimental data. **(G)** Model-derived growth landscapes illustrating the effects of initial fraction of *E. coli* within the community and relative affinity ratios of exploiter and producer (*K*_*s,x2*_ for *C. glutamicum* /*K*_*s,x1*_ for *E. coli*) for the shared resource. Growth rates represent averages until a total cell area of 2,500 µm^2^ is reached. The white line indicates the baseline model assumption.

Tracking species-specific cell area over time revealed that also at this scale *E. coli* and *C. glutamicum* converged to a stable community composition, independent of the initial inoculation ratio (Figure 3E). In the presence of WT *E. coli, C. glutamicum* accounted for approximately 70% of the final population area (*N* = 24 chambers), whereas its fraction remained minimal when enterobactin was absent and *E. coli* Δ*entC* dominated through residual growth (*N* = 27). Although this equilibrium differed from the approximately balanced outcome observed in liquid culture and on plates (Figure 2G-I), all systems consistently showed stable coexistence at a defined community composition rather than unrestricted exploitation.

To determine whether the population size of *C. glutamicum* does affect *E. coli* growth at the microcolony scale due to enterobactin exploitation, growth rates for each organism were calculated between image frames (0.5 h frame rate) across microcolonies of different sizes and compositions (Figure 3F). Whereas *E. coli* growth was largely insensitive to colony composition, *C. glutamicum* growth strongly depended on the relative abundance of exploiter cells and declined sharply once *C. glutamicum* exceeded ∼70% of the colony. This pattern is consistent with the idea that enterobactin remains more readily available to the producer, whereas exploitation by *C. glutamicum* is less effective.

To conceptually and quantitatively explore the social behavior observed in macrocolonies, we implemented a simplified ordinary differential equation (ODE) model that was directly fitted and compared to the experimentally derived single-cell data. To this end, diverging affinities, based on Monod kinetics (kS-values) were assumed to test the possibility of different enterobactin-receptor affinities for both organisms (Figure 3F, G). Although this model does not directly prove the underlying mechanism, it successfully reproduced the experimentally observed interdependence of growth rates when the exploiter was assumed to have 10-fold lower affinity for the shared resource than the producer and consumption and uptake rates were fitted to the given microfluidic data. The model further predicted that increasing enterobactin-receptor affinities of the exploiter promotes collapse of the producer population and can lead to a “tragedy of the commons” [44], whereas coexistence is stabilized when the producer retains superior access to the shared siderophore (Figure 3G). Notably, the model predicts that very high exploiter affinity is also detrimental to the exploiter itself.

To experimentally assess the postulated difference in affinity, we calculated EC_50_ (half-maximal effective concentration [45]) values from growth-rate responses to increasing enterobactin concentrations. This analysis indicated a higher apparent affinity for enterobactin in *E. coli* Δ*entC* than in *C. glutamicum* (EC_50-ΔentC_ = 0.296 ± 0.039 µM; EC_50-Cg_ = 0.744 ± 0.087 µM; Figure S5E, F), thus supporting the model-based hypothesis. Importantly, these values do not directly reflect receptor-siderophore binding affinity, but rather the overall efficiency with which enterobactin exploitation is converted into iron uptake and growth. Together, these findings support our hypothesis in which *E. coli* captures enterobactin more efficiently, likely via its dedicated high-affinity receptor FepA, while *C. glutamicum* accesses only a shared and spatially available fraction of the siderophore. This limits exploitation and stabilizes pairwise coexistence, with community dynamics shaped by the interplay of initial composition, siderophore availability, and species-specific affinity.

### 3.5 Pyoverdine acts as a selfish good in a three species community by preventing *C. glutamicum* enterobactin exploitation

Siderophore-based microbial interactions in natural contexts are more complex than expressed in one-to-one relationships. To address this in our model system, we extended our public good producer/exploiter system toward a multispecies setting. To this end we chose *Pseudomonas putida* as a third interacting organism, which produces the mixed-type siderophore pyoverdine. As expected, pyoverdine could not be utilized by *C. glutamicum*, as shown in pairwise liquid culture and macrocolony assays (Figure 4A, Figure S6B, D-E). Likewise, *E. coli* was also unable to utilize pyoverdine, as inferred from co-cultivation assays with the Δ*entC* mutant (Figure S6C-E). However, *P. putida* efficiently exploited enterobactin produced by *E. coli* – as shown by using the pyoverdine-deficient *P. putida* Δ*pvdD* mutant (Figure S6 and S7). The Δ*pvdD* strain carries a deletion of the gene of the non-ribosomal peptide synthetase PvdD, which is essential for the assembly of the peptide backbone of pyoverdine [46].

**Figure 4:**
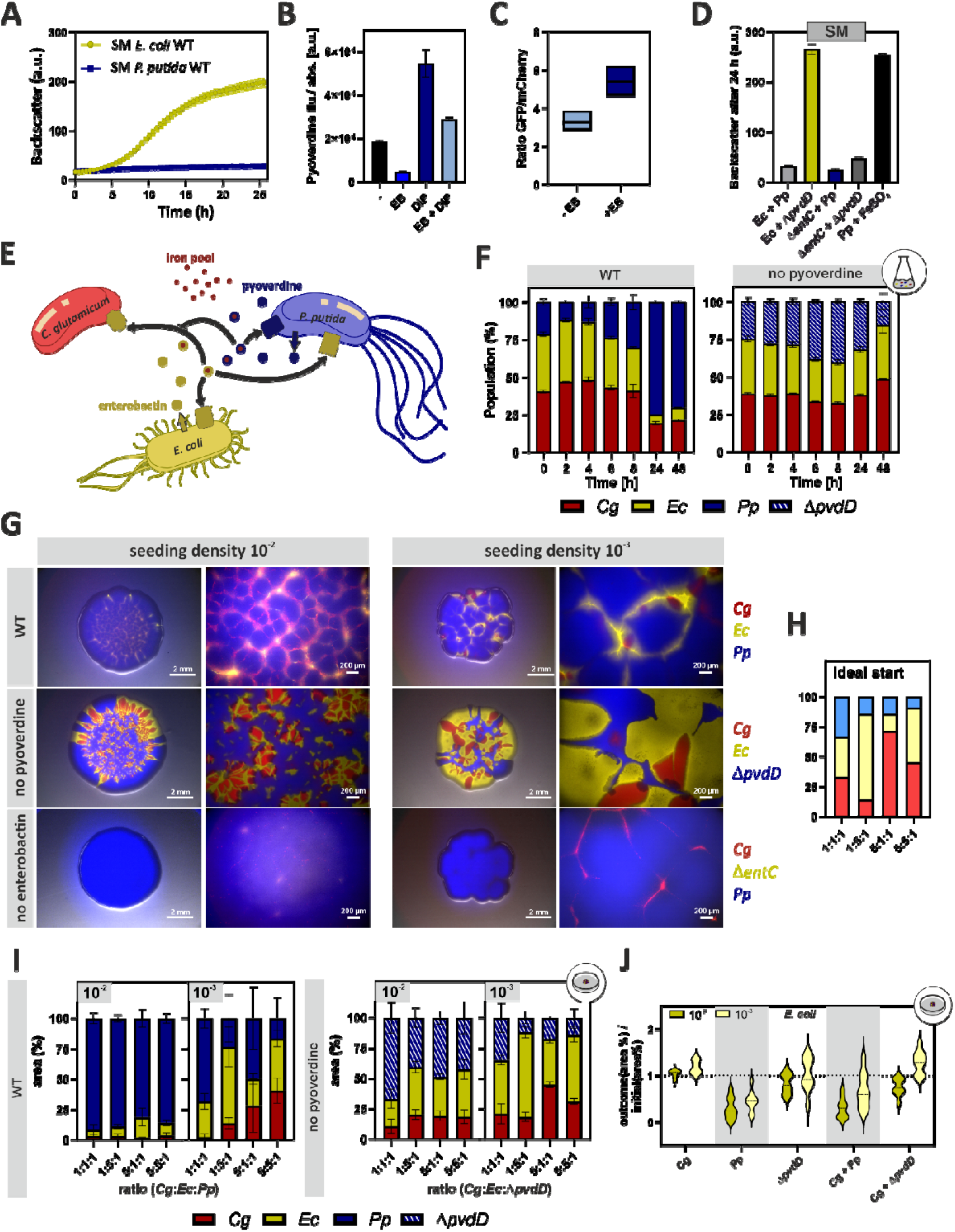
Siderophore-mediated interaction in the trio-consortium of *C. glutamicum, E. coli* and *P. putida* reveals siderophore characteristics as public and selfish goods. **(A)** Growth of *C. glutamicum* with either 50% CGXII-based SM from *E. coli* WT (yellow) or *P. putida* WT (blue). *n* = 3 biological replicates. **(B)** Bar plot of fluorescence measured at λ_ex_ = 398 nm and λ_em_ = 455 nm from *P. putida* cultivation with and without enterobactin (EB) and/or DIP supply, representing pyoverdine levels. **(C)** Comparing iron content by utilizing plasmid-based IronSenseR-mCherry in *P. putida* cultivated with or without 10 µM enterobactin in the presence of 250 µM DIP. **(D)** Bar plot of backscatter (a.u.) at 24 h of *C. glutamicum* on CGXII-based SM mixes of either 30% *E. coli* WT with 20% *P. putida* WT (light grey), 30% *E. coli* WT with 20% *P. putida* Δ*pvdD* (yellow), 30% *E. coli* Δ*entC* with 20% *P. putida* WT (blue), 30% *E. coli* Δ*entC* with 20% *P. putida* Δ*pvdD* (grey) or 50% *P. putida* supplemented with 3.6 µM FeSO_4_ (dark grey). *n* = 3 biological replicates. **(E)** Schematic depiction of the siderophore-mediated interactions within a trio consortium consisting of *C. glutamicum* (public good exploiter), *E. coli* (public good provider) and *P. putida* (selfish good producer and public good exploiter). **(F)** Flow cytometry measurements of co-cultured *E. coli-gfp, C. glutamicum-E2-crimson* and *P. putida-bfp* (left) or Δ*pvdD* (right) under iron limited conditions. *n* = 3 biological replicates. **(G)** Representative stereomicroscopy pictures of mixed drop assay on iron-restricted plates with *C. glutamicum*-*E2-crimson* (red), *E. coli*-*gfp* (yellow) and *P. putida-bfp* (blue), as well as siderophore-deficient mutants, as indicated in two different seeding densities in a 1:1:1 ratio and 16x vs. 80x zoom. **(H)** Schematic overview on ideal start conditions as inoculated for plate assays. **(I)** Quantified area of *C. glutamicum* (red), *E. coli* (yellow) and *P. putida* (blue) deduced from the microscopy pictures as exemplarily shown in (G), set in relation to 100%. *n* = 9 biological replicates. **(J)** *E. coli* outcome analysis with a ratio of *E. coli* outcome over initial area in percentage for two seeding densities (10^-2^ and 10^-3^) in all duo and trio co-cultivations at 1:1:1 ratio.

Enterobactin uptake by *P. putida* was further supported by its physiological response. In the presence of pure enterobactin, and likewise in *E. coli* WT spent medium, *P. putida* produced less pyoverdine (Figure 4B, Figure S7C,D), consistent with reduced intracellular iron limitation. Furthermore, by using the IronSenseR-mCherry variant in *P. putida*, we observed increased intracellular iron levels in the Δ*pvdD* strain upon addition of enterobactin (Figure 4C), confirming that *P. putida* can acquire iron from enterobactin. Previous studies already showed homology between the enterobactin receptors FepA of *E. coli* and PfeA of *P. aeruginosa* [47]. Taken together, the described properties establish *P. putida* as a new consortium member capable of acting as a second enterobactin exploiter producing the non-exploitable (selfish) siderophore pyoverdine.

We next tested how simultaneous exposure to enterobactin and pyoverdine affects *C. glutamicum* growth. To mimic a three-species siderophore environment in monoculture, we combined spent media from *E. coli* WT or Δ*entC* with spent media from *P. putida* WT or Δ*pvdD* (30% and 20%, respectively; Figure 4D). *E. coli* WT spent medium supported growth under iron limitation, but addition of *P. putida* WT spent medium abolished growth despite the presence of enterobactin. Growth was restored with pyoverdine-deficient Δ*pvdD* spent medium as well as by iron supplementation. Thus, pyoverdine can act as a selfish iron-scavenging good that restricts *C. glutamicum* access to iron, even when exploitable enterobactin is present.

### 3.6 Spatial context determines community composition in the three-species consortium

Having shown that pyoverdine can override enterobactin-mediated growth promotion under iron limitation, we next examined how this interaction shapes a three-species consortium across liquid and spatially structured environments.

In liquid microtiter cultivations under iron restriction, using a 1:1:1 inoculation ratio, WT *P. putida* became strongly enriched after 24 h (Figure 4F). Notably, *P. putida* initially declined during the first hours, indicating that its dominance was not immediate but emerged over time. This outcome is consistent with *P. putida* combining exploitation of enterobactin with pyoverdine production, which limits iron access for the other community members. Accordingly, when pyoverdine production was abolished, the community remained close to the inoculated ratio after 24 h. Conversely, in the absence of enterobactin, WT *P. putida* prevailed, whereas Δ*pvdD*-containing combinations showed little to no growth (Figure S8C,D). Under iron-replete conditions without chelator, *P. putida* did not become overrepresented (Figure S8A,B), confirming that its dominance depends on iron limitation.

We next asked whether spatial structure modifies this interaction by restricting siderophore diffusion and access. On iron-limited CGXII plates, WT *P. putida* again displaced both partners at high seeding density, while *C. glutamicum* and *E. coli* persisted at lower abundance than in liquid culture (10^−2^; Figure 4G, Figure S9). However, spatial structure also enabled local niche formation. In the Δ*pvdD* background, *E. coli* and *C. glutamicum* grew in close proximity, whereas *P. putida* Δ*pvdD* was largely confined to peripheral regions. As expected, loss of enterobactin production in *E. coli* enabled *P. putida* to dominate the community. Thus, when diffusion is limited, the balance between public enterobactin and selfish pyoverdine becomes locally structured rather than uniformly shared across the consortium.

Varying inoculation ratio and seeding density further revealed strong context dependency (Figure 4G–I, Figure S10). Under 1:1:1 WT conditions, *P. putida* became overabundant at the high inoculation density, whereas the lower density and higher initial *E. coli* abundance improved its persistence, consistent with increased spatial separation and reduced siderophore piracy. At 10^−3^ seeding density, *C. glutamicum* grew in distinct patches surrounded by *E. coli*, separating it from *P. putida*. Without pyoverdine, *E. coli* and *C. glutamicum* benefited from high seeding density, whereas lower density maintained compositions closer to the inoculated ratios. Without enterobactin, *P. putida* remained dominant across conditions, whereas under iron-replete conditions *C. glutamicum* outcompeted both partners (Figure S10).

We finally asked the question of what is the impact of exploitation on the producer. We therefore quantified final-to-initial area ratios and demonstrated that *P. putida* significantly impaired *E. coli* growth despite the higher affinity of enterobactin according to literature values [9] (Figure 4J). This effect was strongly reduced in the Δ*pvdD* background, although its magnitude varied with initial conditions. Overall, *P. putida* restricts *E. coli* growth while consistently benefiting from co-cultivation with *E. coli*, particularly when *C. glutamicum* is also present (Figure S10F). By contrast, *C. glutamicum* exploitation did not affect *E. coli* growth, consistent with our previous results (Figure 3F).

To further resolve how initial community composition influences community dynamics, we performed analogous experiments in microfluidic chambers and followed microcolony development, composition and architecture by time-lapse microscopy (Table S2, Figure 5). Ternary plots revealed that high initial abundances of either *C. glutamicum* or *P. putida* resulted in a pronounced shift towards *P. putida* dominance, often reaching 70–100% of the final community. By contrast, when *E. coli* initially represented more than 50% of the community, it persisted at high abundance, approximately 80–90%, while also supporting *C. glutamicum* growth (Figure 5A, Videos S3–5). Replacing WT *P. putida* with the Δ*pvdD* mutant reduced *P. putida* overabundance, with its final proportion stabilizing at approximately 50% (Figure 5B, Videos S6–8). Notably, *C. glutamicum* also grew in some chambers in the presence of WT *P. putida*, but only when the initial spatial arrangement allowed *C. glutamicum* patches to form within surrounding *E. coli* cells, thereby locally separating it from *P. putida* (Video S3).

**Figure 5:**
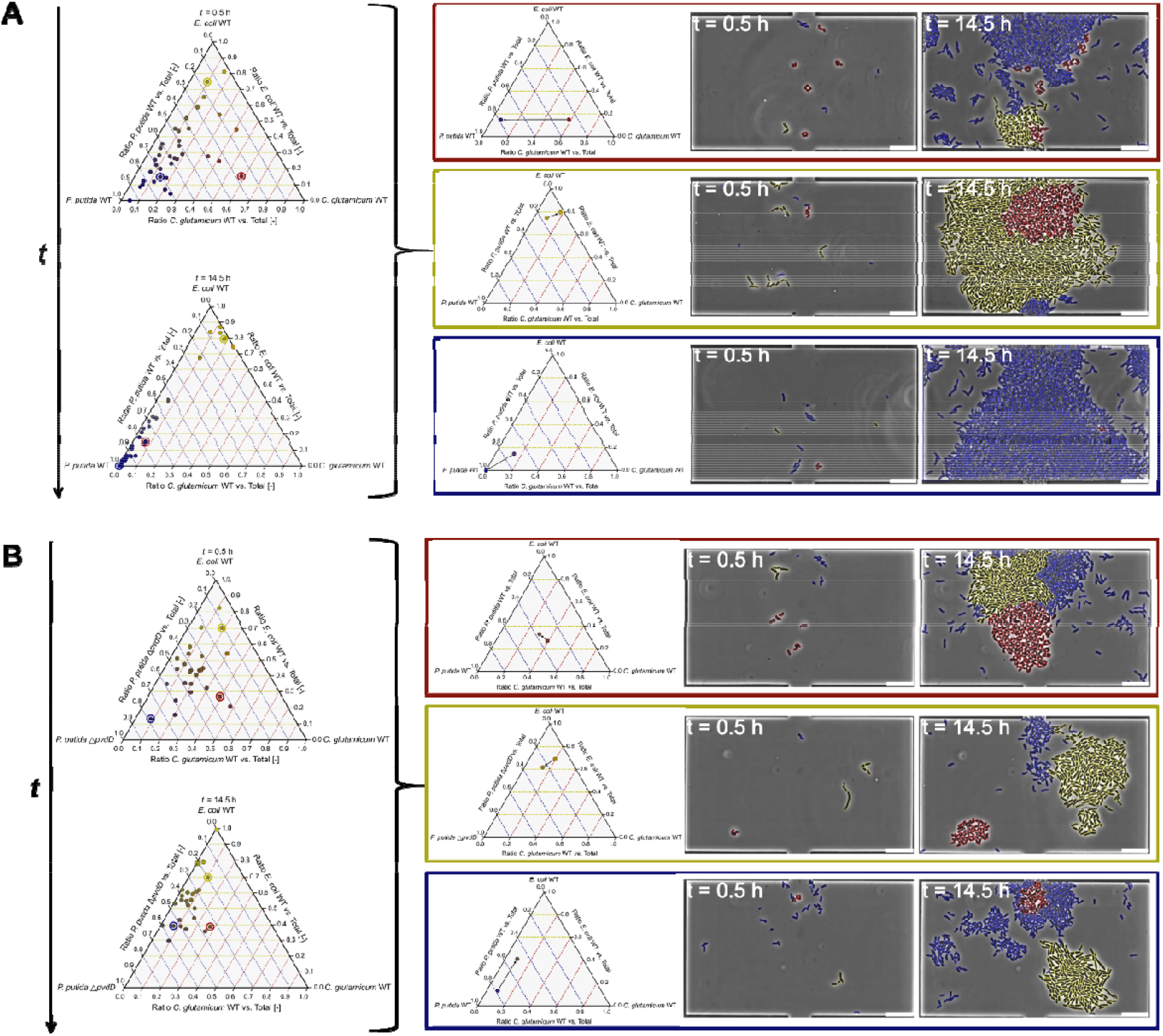
Microfluidic single-cell analyses of community composition dynamics in three-species consortia. **(A)** Ternary plots showing the relative abundance of *C. glutamicum* WT (red), *E. coli* WT (yellow) and *P. putida* WT (blue). Each point represents a single chamber, positioned according to the fractional cell area of each species. The plots show the initial composition at 0.5 h (top) and the composition after 14.5 h (bottom). Chambers with strong dominance of one species are highlighted by colored outlines and corresponding representative chambers are shown to the right, including their temporal development and segmented images at *t* = 0.5 h and 14.5 h. **(B)** As in (A), but with *P. putida* Δ*pvdD* instead of WT. All experiments were performed in CGXII medium with 3.6 µM FeSO_4_ at 30 °C. All scale bars 10 µm.

Collectively, these findings show that siderophore-mediated interactions in multispecies communities are highly context dependent. In this three-species community, enterobactin produced by *E. coli* functions as a public good for both *C. glutamicum* and *P. putida*, whereas pyoverdine acts as a selfish good by reducing iron availability for competitors lacking compatible uptake systems. Spatial organization and initial community composition strongly modulate these interactions by determining the extent to which diffusible siderophores are shared, recaptured, or rendered inaccessible.

## Discussion

Over the past two decades, siderophore-mediated social interactions have largely been studied in pairwise systems, revealing that bacteria can adopt distinct ecological roles as producers, cheaters or exploiters. Producers synthesize and secrete siderophores, cheaters benefit from siderophores produced by clonemates, and exploiters access xenosiderophores released by other species [2,22,48-50]. However, natural microbial communities are not built from isolated pairwise interactions. They contain multiple species with different siderophore repertoires, uptake systems, affinities and spatial arrangements. Consequently, the ecological function of a siderophore is not fixed, but emerges from the community context in which it is produced and used [22,23,27,51]. In this study, we demonstrate that enterobactin from *E. coli* acts as a public good that can support the growth of the exploiters *C. glutamicum* and *P. putida* under iron limitation. On the other hand, pyoverdine produced by *P. putida* acts as a selfish good by inhibiting growth of the competing community members. These results support a broader view of siderophores as context-dependent iron-allocation agents that can either promote coexistence or enforce exclusion, depending on the surrounding community and physical environment.

Corynebacteria are increasingly recognized as common siderophore exploiters in natural microbiomes, yet the extent and ecological consequences of this strategy have remained difficult to dissect mechanistically [52]. By using *C. glutamicum* as a genetically tractable model organism, we demonstrate that a non-producing bacterium can exploit chemically diverse xenosiderophores from bacteria and fungi. Such diversity is consistent with adaptation to iron-limited, siderophore-rich environments and raises a broader evolutionary question: Under which ecological conditions it is more advantageous to maintain uptake capacity for diverse exogenous siderophores than to invest in siderophore biosynthesis? One potential hypothesis is that, in communities with public good producers, exploitation is metabolically less costly than de novo synthesis, consistent with the logic of the Black Queen hypothesis, whereby organisms can benefit from losing costly functions, if these are provided by other members of the community [53,54]. This logic is mirrored in *Pseudomonas* communities, where cheaters can persist, if the producer population is stable and sufficiently high [23,55].

Remarkably, enterobactin exploitation by *C. glutamicum* is constrained. Genetic evidence, metabolite measurements, and biosensor readouts all support direct uptake of enterobactin-derived iron rather than an indirect growth-promoting effect. At the same time, across structured plates and microfluidic chambers, exploitation never resulted in a collapse of the producer population. Instead, *C. glutamicum*-*E. coli* co-cultivation reproducibly converged to stable community compositions. Modelling and EC_50_ analyses further suggest that this stability may be related to asymmetric access to enterobactin. The producer, carrying the cognate FepA receptor for uptake, may capture enterobactin more efficiently than the exploiter. In this scenario, the exploiter accesses only the fraction of enterobactin that escapes producer recapture. This suggests a simple ecological principle, namely that stable interspecies exploitation of a diffusible public good may require asymmetry in access, such that the producer retains privileged uptake. This interpretation is consistent with previous work, showing that enterobactin can exhibit density-dependent privatization, remaining partially cell-associated at low cell densities and becoming increasingly shared as extracellular enterobactin accumulates [56]. If exploiter public good access becomes too efficient, the shared resource may no longer stabilize coexistence but instead drives a “tragedy of the commons” scenario as is often observed during interactions between producers and cheaters. In this scenario, cheaters can still take up the corresponding siderophore-iron complexes via the same receptors and thus with the same efficiency as the producers [2,44,51]. Although this is not observed during interaction between *C. glutamicum* and *E. coli*, a related stabilizing principle has been proposed for cross-feeding mutualisms. When exploiter and producer have similar uptake affinities, stable coexistence may still be maintained if the exploiter reciprocates by providing a public good or another trait of comparable ecological value [57-59]. Exemplarily, this was recently observed in *Burkholderia contaminans* making use of the siderophore bacillibactin produced by *Bacillus velezensis*, while providing branched-chain amino acids [35].

The three-species community revealed an additional layer of ecological complexity. Although *P. putida* can exploit enterobactin, its own pyoverdine was inaccessible to both *C. glutamicum* and *E. coli*. As a result, pyoverdine abolished the growth benefit that enterobactin otherwise provided to the non-producing exploiter. This is especially interesting considering enterobactin has been reported to possess a higher binding affinity to iron (pFe ≈ 34.3) [60,61] than pyoverdine (pFe ≈ 27) [62]. Thus, our observation revealed that the ecological outcome of multispecies communities cannot be predicted solely from siderophore binding affinities. Instead, it depends on how efficient the iron-siderophore complex can be accessed by individual species. This mechanism is certainly relevant beyond the here described model consortium, particularly in communities containing pyoverdine-producing pseudomonads and taxa with only partially overlapping siderophore networks [23,63].

On plates, changes in seeding density and inoculation ratio shifted community outcomes. Higher spatial segregation at lower seeding densities reduced the access of *C. glutamicum* to diffusible enterobactin and partially buffered the dominance of *P. putida*. In well-mixed cultures, the dominance of *P. putida* emerged more slowly, with an initial decline of *P. putida* in the early stages, suggesting that homogenous enterobactin distribution promotes early growth of *C. glutamicum* and *E. coli*. Together, these observations show that the same siderophore network can produce different ecological outcomes depending on spatial arrangement and starting community composition [64]. The molecules remain the same, but the accessibility landscape changes.

In summary, our results shift siderophore-mediated interactions from fixed categories of cooperation, cheating or exploitation toward a dynamic framework of iron allocation. Using *C. glutamicum* as a model non-producing exploiter, we show that xenosiderophore use can be constrained, allowing access to a shared siderophore without destabilizing the producer population. In multispecies communities, this balance can be reshaped by additional siderophores, such as pyoverdine, that restrict iron access for competitors and thereby redistribute benefits across the community [65-67]. More broadly, our findings suggest that siderophore chemistry, uptake compatibility, relative affinity, and spatial structure together determine who benefits from different iron-siderophore complexes and who is excluded from it. This framework should be useful for understanding iron competition in natural microbiomes and for designing siderophore-shaped synthetic communities in biotechnology, agriculture [68,69], and host-associated systems [65].

## Material and Methods

### Microbial strains and cultivation conditions

All microbial strains used in this study are listed in Table S3. Microbial strains were streaked from frozen 20% glycerol stocks on 1.5% (w/v) agar medium containing complex medium. For *C. glutamicum*, brain heart infusion (BHI) (Difco, BD, Heidelberg, Germany) (37 g/L BHI) medium was used as complete medium. All other microbial strains were cultivated in lysogeny broth (LB) (10 g/L tryptone, 5 g/L yeast extract, 10 g/L NaCl) medium, while *Ustilago maydis* was cultivated in LB or yeast extract-peptone-sucrose (YEPS) (10 g/L yeast extract, 10 g/L peptone, 10 g/L sucrose) medium. Strains were cultivated exclusively at 30°C, with exception for *E. coli* in first pre-cultures at 37°C. For liquid cultivations, single colonies from agar plates were picked and incubated in 5 ml complex medium in reaction tubes at 170 rpm or in 1 ml medium in deep-well plates (VWR International, Radnor, PA, United States) at 900 rpm. Upon cultivating for approximately 8 h in BHI or LB medium, cells were diluted 1/10 and used to inoculate a second preculture in CGXII minimal medium (20 g/L (NH_4_)_2_SO_4_, 1 g/L K_2_HPO_4_, 1 g/L KH_2_PO_4_, 5 g/L urea, 42 g/L 3-morpholinopropane-1-sulfonic acid (MOPS), 13.3 mg/L CaCl_2_*2H_2_O, 0.25 g/L MgSO_4_*7H_2_O, 0.2 mg/L biotin, 0, 1 or 10 mg/L FeSO_4_*7H_2_O, 10 mg/L MnSO_4_*H_2_O, 0.1 mg/L ZnSO_4_*7H_2_O, 0.02 mg/L CuSO_4_*5H_2_O, 0.002 mg/L NiCl_2_*6H_2_O; pH was adjusted with KOH to pH of 7.0.) [70] containing 2% (w/v) glucose as carbon source. To starve cells of the second pre-culture from iron prior to experiments, CGXII minimal medium without FeSO_4_ was used (referred to ‘iron-free CGXII’). Note that protocatechuate (PCA) (3,4-dihydroxybenzoic acid), a ferric chelator, is usually added to the CGXII minimal medium to facilitate iron uptake and was therefore left out in our cultivations [70,71]. For *E. coli*, 1 µM FeSO_4_ was added to the second pre-culture. Second pre-cultures were performed in 20 - 25 mL minimal medium in shaking flasks at 120 rpm or in 1 mL medium in 96-deep-well plates (VWR International, Radnor, PA, United States) at 900 rpm.

Main cultures were inoculated at an optical density at 600 nm (OD_600_) of 1, if not stated otherwise, and either performed in shaking flasks in respective volumes or for online monitoring of growth in 750 µl in 48-well microtiter Flower Plates in a BioLector I microtiter cultivation system (Beckman Coulter GmbH, Aachen, Germany) [72]. In the microtiter cultivation system, backscatter (a.u.) was measured in 30 min intervals as scattered light at *λ*: 620 nm (signal gain: 20). *P. putida* cultures were inoculated at an OD_600_ of 0.5 in a volume of 1ml. As standard conditions, CGXII minimal media was used as described above. For analysis of growth in presence of LB-based spent media, cells were inoculated in 2-fold concentrated iron-free CGXII medium supplemented with 2*2% glucose and 2*150 µM 2,2′-dipyridyl (hereafter referred to as DIP). For CGXII-based spent media, cells were inoculated in 2-fold concentrated CGXII medium with 2*2% glucose, 3.6 µM FeSO_4_ and no DIP. Each cell-media suspension was mixed with 50% of respective spent medium. Note: The addition of DIP is a well-established strategy to mimic non-accessible iron pools in siderophore studies [73-76]. Siderophores have a higher affinity for iron than this chelator, thereby enabling iron uptake by those cells, which can use the specific siderophore. When enterobactin was added as pure compound (Sigma-Aldrich, dissolved in DMSO), 3.6 µM FeSO_4_ and 150 µM DIP was added in liquid cultivations.

Bacterial strains containing plasmids mediating a kanamycin resistance were cultivated in the presence of appropriate amounts of the antibiotic (*C. glutamicum*: 25 µg/L; *E. coli*: 50 µg/L) to ensure plasmid maintenance. If needed, LacI-P_*tac*_-derived gene expression was induced by the addition of isopropyl-β-D-1-thiogalactopyranoside (IPTG).

### Production and harvesting of siderophore containing spent medium

Spent medium refers to the siderophore-enriched growth medium remaining after cultivating of a respective siderophore-producing organism, but cells removed.

For screening purposes, LB-based spent medium was used: Cells of siderophore-producing strains from an overnight culture were transferred at an OD_600_ of 0.45 to 50 mL LB medium supplemented with 200 µM DIP, while *U. maydis* cells were cultivated in YEPS medium containing 300 µM DIP.

For more specific analyses, CGXII-based spent medium was used: *E. coli* or *P. putida* strains from an overnight culture were transferred at an OD_600_ of 0.45 to 50 mL CGXII media as described above, but containing 0.36 µM FeSO_4_ and 360 µM DIP.

After two days of cultivation in shake flasks, the culture supernatant was harvested and sterile filtered (0.2 µm). Harvested spent medium was stored at −20°C until further use.

### Recombinant DNA work

Standard molecular methods, such as DNA restriction, PCR, and Gibson assembly, were performed according to the manufacturer’s instructions or to previously described standard protocols [77,78]. All plasmids and primers used in this study are listed in Tables S4 and S5. Oligonucleotide synthesis and Sanger DNA sequencing was performed by the company Eurofins Genomics (Ebersberg, Germany).

In this work, all plasmids were constructed using the enzymatic Gibson assembly approach. Plasmids were transferred into competent *C. glutamicum* cells by electroporation as described previously [79]. For genomic deletions and integrations, derivates of the suicide vector pK19-*mobsacB* were used [80]. Two step homologous recombination and selection was done as described previously [81].

Plasmids were transferred into competent *E. coli* cells by heat shock or TSS transformation [82,83]. Genomic deletion of the *entC* gene in strain *E. coli* PC 0886 Δ*entC* was achieved with the Lamda red recombinase-based deletion method described previously [84]. The disruption cassette consisted of the *neoR* resistance gene, encoding the neomycin phosphotransferase providing resistance against neomycin and kanamycin. This gene was flanked by FLP recombinase recognition target (frt) sites, which are needed for later elimination of the *neoR* gene. The up- and downstream sequences of *entC* (approximately 200 nt) formed the ends of the disruption cassette long sequences, allowing for replacement of the *entC* gene by the disruption cassette by homologous recombination.

For genomic integrations, the deletion protocol was modified. An integration cassette was used instead of the disruption cassette. The design was comparable to the disruption cassette, but the sequence that should be integrated was inserted between the *neoR* – frt sequence and the right homologous flank.

For the genomic integration of the constitutive mTagBFP (for fluorescent labeling) and the IronSenseR-mCherry, tetra parental patch-mating was performed on 1% LB agar plates overnight at 30 °C. Freshly streaked strains of *P. putida* KT2440 WT or *P. putida ΔpvdD* were plated as acceptor strains. Helper strain *E. coli* HB101 pRK2013 and *E. coli* DH5αλpir pTNS1 either helped with pili formation or delivered necessary transposase genes respectively. Lastly a *E. coli* Pir2 carrying the miniTn7 plasmid with the IronSenseR-mCherry or constitutively expressed fluorescent marker gene (*mTagBFP*) was used as a donor strain. After incubation, cells were streaked on LB plates containing gentamycin (25 μg mL^−1^) and irgansan (25 μg mL^−1^) and cultivated over night to select for *P. putida* KT2440 WT or *P. putida ΔpvdD* integration cassette. Then single clones were streaked on LB containing kanamycin (25 μg mL^−1^) and irgansan (25 μg mL^−1^) as well as LB containing gentamycin (25 μg mL^−1^) and irgansan (25 μg mL^−1^). Clones only growing on LB containing gentamycin and irgansan were likely carrying the Tn7 integration, which was then confirmed *via* colony PCR.

### Siderophore sharing in structural environments

Double layer agar plate assays were performed to study siderophore sharing in structural environments. LB or CGXII plates consisted of a solid 1.5% agar bottom layer and a soft 0.4% agar top layer to facilitate diffusion. Plates were prepared both with and without DIP to compare the effects of ‘free iron’ availability versus a condition where iron could only be acquired via siderophores. For LB-based agar plates, 200 µM DIP was added, but no additional iron. For CGXII-based agar plates, 3.6 µM FeSO_4_ was added and 100 µM DIP was utilized, while for ‘full-iron conditions’ 36 µM FeSO_4_ were used without any addition of DIP.

#### Xenosiderophore utilization screen (LB-based)

Cells from LB overnight cultures of different putative siderophore producers (Table S3) were transferred into 0.9% NaCl at cell density of 0.24*10^10^ cells/ml. 40 µL cell suspension were streaked on freshly prepared plates as thick lane. After overnight cultivation of producer strains at 30°C, iron-starved *C. glutamicum* cells were transferred into 0.9% NaCl at cell densities of 10^10^, 10^9^, 10^8^, 10^7^, 10^6^, 10^5^ and 10^4^ cells/mL. 3 µL of these cell suspensions were dropped at different distances to the lane of siderophore producers. Images were taken after 48 h cultivation at 30°C.

#### Siderophore-mediated co-cultivations (CGXII-based)

Cells were cultivated in iron-free CGXII as second pre-culture. Subsequently, cells corresponding to an OD_600_ of 1 were washed and resuspended in 0.9% NaCl. Each 2 µL of cell suspension was dropped on the CGXII-soft agar layer either as monocultures, or in duo or trio combinations dropped ∼ 5 mm apart. Plates were incubated at 30 °C for 3 days and images were taken under a steromicroscope Zeiss Axio Zoom V16 with respective filter units (GFP: *λ*_Ex_: 489 nm, λ_Em_: 509 nm (*gfp*); HE mPlum: λ_Ex_: 587 nm, λ_Em_: 649 nm (*E2-crimson*); DAPI: λ_Ex_: 358 nm, λ_Em_: 463 nm (*bfp*)) once after 48 h and 120 h. Analysis of percentage area was performed using the open source program ImageJ/Fiji [85,86]. Area per fluorescent channel image was extracted using individual background correction thresholds.

### *In vivo* measurement of the biosensor for iron in *C. glutamicum*

For the *in vivo* online measuring iron levels in *C. glutamicum*, the fluorescent protein LSSmApple of the original IronSenseR [42] was replaced by mCherry, constructing IronSenseR-mCherry plasmid. For *C. glutamicum*, an N-terminal deca-histidine tag followed by a TEV cleavage site was directly fused to the biosensor encoding gene cassette. Electrocompetent *C. glutamicum* were transformed with the plasmid pPREx210xHis-TEV-MDtxR_G149_GC. Single colonies were cultivated in 800 µl ml BHI supplemented with kanamycin in a deep-well plate at 30 °C and 900 rpm for 8 h. The second pre-culture was prepared in iron-free CGXII medium containing 2% (w/v) glucose supplemented with 100 µM IPTG. Cultures were incubated overnight (16 h) at 30 °C and 900 rpm in deep-well plates. For the main culture, cells were inoculated to an initial OD_600_ of 0.5 in 10 mL CGXII minimal medium supplemented with 2% (w/v) glucose, 3.6 µM FeSO_4_ as iron source, no PCA and 100 µM IPTG. Where indicated, enterobactin was added to a final concentration of 10 µM. Cultures were incubated in 100 mL shaking flasks at 30°C for 2 h on a rotary shaker. Cells corresponding to an OD_600_ of 0.5 in 1 mL culture volume were harvested by centrifugation (5,000 rpm, 4°C, 5 min), washed, and resuspended with ice-cold MOPS buffer (20 mM, pH 7.0). A volume of 100 µL was transferred to a microplate and analyzed using a multimode microplate reader (Infinite M1000Pro, Tecan). Emission spectra were recorded in 5 nm increments from 490 to 750 nm for GFP upon excitation at 453 nm, and from 580 to 750 nm for mCherry upon excitation at 550 nm. Fluorescence values were corrected by subtracting the signal obtained from buffer-only controls. To obtain the ratios, the peak of GFP fluorescence (λ_em_= 510 nm) was divided by the peak value of mCherry (λ_em_= 610 nm). For *P. putida* the strain carrying the integration of the IPTG inducible IronSenseR *(P. putida ΔpvdD ::attTn7 IronSenseR-mCherry*), was inoculated in LB at 30 °C and 1200 rpm overnight. The second preculture was inoculated at an OD_600_ of 0.5 and incubated overnight in LB at 30 °C and 1200 rpm with the addition of 500 µM IPTG. For the main culture, cells were inoculated to an initial OD_600_ of 0.1 in 1 mL LB 500 µM IPTG. Where indicated, 10 µM enterobactin and 250 µM DIP was added. Cultures were incubated for 6 hours at at 30 °C and 1200 rpm. 100 µL of the culture was harvested washed and resuspended in 100 µL PBS, transferred to a microtiter plate and analyzed using a multimode microplate reader (Infinite M1000Pro, Tecan). Two distinct emission spectra of GFP (λ_ex_ = 480 nm, λ_em_ = 500-600 nm) and mCherry (λ_ex_ = 587 nm, λ_em_ = 600-700 nm) were recorded. To obtain the ratios, the peak of GFP fluorescence (λ_em_ = 508 nm) was divided by the peak of mCherry (λ_em_ = 608 nm).

### MS-measurements of enterobactin and organic acids

For MS measurements, four replicates of *C. glutamicum* were cultivated as described above in an iron-free CGXII pre-culture. In the main culture, *C. glutamicum* was cultivated at a starting OD_600_ of 1 in 50 mL CGXII supplemented with 2% glucose and either (i) 3.6 µM FeSO_4_, 150 µM DIP, no PCA (iron limited) (ii) 3.6 µM FeSO_4_, 10 µM enterobactin, 150 µM DIP, no PCA or (iii) 36 µM FeSO_4_. Samples were taken after 5, 10, 30, 120, 300 and 1440 minutes.

For measurements of extracellular enterobactin, cells were centrifuged and supernatant filtered (0.22 µm). For intracellular measurements of enterobactin and organic acids, cells corresponding to 2 OD units were washed in 10 mL 0.9% NaCl and cell culture was rapidly filtered through a nylon membrane with a pore size of 0.45 µm using a vacuum filtration system (Merck Millipore). The filter was washed twice with 5 mL ice-cold 0.9% NaCl. The membrane was immediately placed into 2 mL tubes preloaded with steel (2 mm) and glass beads (Retsch) and snap-frozen in liquid nitrogen. The entire sampling and quenching procedures were completed within 45 seconds.

Enterobactin and organic acids were extracted from the cells using a method adapted from [87]. For extraction, 1 mL of solvent consisting of acetonitrile, methanol, and 0.1 M formic acid (40:40:20, v/v) was added to reaction tubes containing nylon filters with collected cells. The extraction solvent was supplemented with thio-alpha-ATP (TriLink Biotechnologies, 0.5 µM in 10 mL), d8-valine and d5-phenylalanine (Cambridge Isotope, each 0.5 µM in 10 mL extraction buffer), which served as internal standards. Cell disruption was achieved using a bead mill (Retsch) operated at 30 Hz for 2 minutes with pre-chilled holders to facilitate cell detachment from the filters and ensure efficient lysis. The resulting extract was neutralized by adding 90 µL of 15% (w/v) ammonium hydroxide.

After removal of the nylon filters and steel beads, samples were incubated at −20 °C for 30 minutes to improve protein precipitation. The samples were then centrifuged at 16,000 × g for 10 minutes at 4 °C, and the clarified supernatant was transferred to a 15 mL tube and diluted with 5 mL of ice-cold water. The diluted extracts were frozen at −80 °C and subsequently lyophilized for 48 hours using an Alpha 1–2 LDplus freeze dryer (CHRIST). Dried samples were stored at −80 °C and reconstituted in 200 µL of water prior to analysis.

For analysis of extracellular enterobactin, culture media was filtrated through a centrifugal filter with a size exclusion of 0.2 µm (VWR) and the cleared extract was immediately frozen in liquid nitrogen. For purification of enterobactin from the culture media, the filtrate was further extracted using solid-phase extraction (Strata-X polymeric reverse phase, Phenomenex) in accordance with the manufacturer’s instructions. d8-valine and d5-phenylalanine (Cambridge Isotope) was added to the culture medium prior SPE as internal standard to a final concentration of 2.5 µM each. SPE eluate was transferred to a 15 mL tube and diluted with 5 mL of ice-cold water. The diluted eluates were frozen at −80 °C and subsequently lyophilized for 48 hours using an Alpha 1–2 LDplus freeze dryer (CHRIST). Dried samples were stored at −80 °C and reconstituted in 200 µL of water prior to analysis.

Enterobactin measurements were performed on an ACQUITY Premier reverse phase UPLC system (Waters) coupled to a Q Exactive Plus quadrupole–Orbitrap mass spectrometer (Thermo Scientific). An injection volume of 5 to 10 µL was introduced into the reverse phase UPLC system. Compounds were separated on an ACQUITY UPLC HSS T3 analytical column (1.8 µm particle size VanGuard FIT, 2.1 mm x 150 mm, Waters) and achieved by generation of a solvent gradient (flow rate: 0.300 ml/min, 0-0.5 min 100% A, 0.5-6 min 100-89% A, 6-8 min 89-0% A, 8-11 min 0% A, 11-12 min 0-100%, 12-17 min 100% A). Solvent A consists of 10 mM Ammonium formiate in 0.15% formic acid and solvent B of 0.1% formic acid in acetonitrile. The column temperature was maintained at 35 °C. Ionization was performed in negative electrospray ionization (ESI) mode with the following source settings: sheath gas 30, auxiliary gas 15, sweep gas 2, spray voltage 3.2 kV, capillary temperature 230 °C, S-Lens RF level 45, and auxiliary gas heater temperature 280 °C. Full-scan mass spectra were acquired over an m/z range of 150–2000 at a resolution of 70,000. The automatic gain control target was set to 1 × 10^6^ ions, with a maximum injection time of 200 ms. Targeted data processing was carried out using Skyline software (version 25.1, MacCoss Lab, University of Washington). Enterobactin was identified as a singly negatively charged species (m/z 668.1369) by high-resolution mass spectrometry, using a reference standard (Sigma-Aldrich, E3910). Enterobactin peak areas were normalized to the internal standards and to OD_600_ values recorded at the time of sampling.

Organic acids were analysed using a Dionex ICS-6000 HPIC system coupled to a Thermo Scientific Q Exactive Plus quadrupole–Orbitrap mass spectrometer, as described by [88]. A 10 µL aliquot of the extracted sample was injected into the IC system using a Dionex AS-AP autosampler operated in push partial mode.

Anion exchange chromatography was performed using a Dionex IonPac AS11-HC analytical column (2 mm × 250 mm, 4 µm particle size; Thermo Fisher Scientific) equipped with a Dionex IonPac AG11-HC guard column (2 mm × 50 mm, 4 µm; Thermo Fisher Scientific) and operated at a column temperature of 30 °C. The mobile phase was generated using an eluent generator with a potassium hydroxide cartridge, producing a potassium hydroxide gradient. The flow rate was set to 380 µL min^−1^, starting at a KOH concentration of 5 mM for 1 min, followed by a linear increase to 85 mM over 35 min, which was held for 5 min. The concentration was then immediately reduced to 5 mM, and the system was equilibrated for 10 min.

Spray stability of aqueous solution was improved using a makeup flow of methanol containing 10 mM acetic acid, delivered by an AXP pump with a flow rate of 150 µL min^−1^. Electrospray ionisation was performed in the ESI source using the following parameters: sheath gas 30, auxiliary gas 15, sweep gas 0, spray voltage −2.8 kV, capillary temperature 300 °C, S-Lens RF level 45, and auxiliary gas heater temperature 380 °C.

Full-scan spectra were acquired over an m/z range of 60–800 at a resolution of 140,000, with an automatic gain control (AGC) target of 1 × 10^6^ ions and a maximum injection time of 500 ms. Organic acids were identified according their retention time as a singly negatively charged species (fumaric acid m/z = 115.0037, citric and isocitric acid m/z 191.0197, succinic acid m/z = 117.0193, aconitic acid m/z = 173.0092, malic acid m/z = 133.0143, oxoglutaric acid m/z = 145.0143) by high-resolution mass spectrometry and using reference standards obtained from Sigma-Aldrich.

Targeted data evaluation was performed using Skyline (version 25.1, MacCos Lab Software, University of Washington). Peak areas were normalized to the internal standards and to OD_600_ values recorded at the time of sampling.

### Flow Cytometry of trio cultures

*P. putida*, and *E. coli* were precultured in LB, while *C. glutamicum* was precultured in BHIS overnight at at 30 °C and 1200 rpm. The second preculture was inoculated at an OD_600_ in CGXII medium with 2% glucose and 3.6 µM FeSO_4_. For the main culture, the three different strains were mixed at similar ratios to a total OD_600_ of 1, in CGXII medium with 2% glucose containing either 0.36 µM FeSO_4_, 3.6 µM FeSO_4_, 0.36 µM FeSO_4_+ 250 µM DIP or 3.6 µM FeSO_4_+ 250 µM DIP and cultivated for 48 hours at 30 °C and 1200 rpm. During cultivation and at the end 20 µL samples were taken washed and diluted to an OD_600_ of 0.05 in 200 µL of PBS. Afterwards samples were analyzed in the flowcytometer (Amnis® CellStreamTM System, Merck, now Cytek Biosciences, California, USA). Cell populations were gated for size and complexity via front scatter (FSC) and side scatter (SSC). Afterwards, populations were further gated for mCherry, GFP or mTagBFP fluorescence. For analysis of the Flowcytometry data, the CellStream Analysis Software (Merck, now Cytek Biosciences, California, USA) was used.

### Pyoverdine measurements

Precultures were performed in LB at 30 °C and 1200 rpm. Main cultures were inoculated at an OD_600_ of 0.05 in LB, with 250 µM DIP, 250 µM DIP and 10 µM enterobactin or no addition. To measure pyoverdine production, after cultivation 20 µl of culture were transferred to 96-well clear bottom microtiter plates and diluted with 80 µL of PBS. Using a plate reader, Infinite M1000 PRO (Tecan Austria GmbH) absorption was measured at 600 nm and pyoverdine fluorescence (λ_ex_ = 398 nm, λ_em_ = 455 nm) was measured in the same device.

### Single-cell cultivation in microfluidic chips

Microfluidic chips were fabricated using soft lithography with a two-layer SU-8 structure on a four-inch silicon wafer. The photomask, devised with Clewin5, delineated cultivation chambers measuring 60 × 100 × 0.9 µm with two opposing 10 µm wide inlets linked to the inlet and outlet of the chip via 9.8 µm deep channels. Each inlet supplied eight rows of thirty chambers each. PDMS (Sylgard 184; base with 10 wt% curing agent) was poured onto the SU-8 master, then cured for 2 h at 80 °C. It was then cut to size and punched to create in- and outlet openings (Ø 0.5 mm). The PDMS plates were bonded to glass substrates (D263® Bio, 39.5 × 34.5 × 0.175 mm) using O_2_ plasma activation (25 s, 510 W, Femto Plasma Cleaner, Diener Electronics, Germany), which enabled covalent bonding via condensation.

All experiments were performed in PCA-free CGXII minimal medium. *C. glutamicum* was pre-cultured at 30 °C in three steps: 8 h BHI, followed by 16 h and 4 h iron-limited CGXII. *E. coli* was grown in LB (37 °C), then 16 h iron-limited CGXII, and finally 4 h CGXII with 3.6 µM FeSO_4_. All cultures were initiated from single colonies on agar plates. CGXII composition followed standard formulations as described above, but adding only 1% (w/v) glucose. Moreover, the addition of DIP was not required to provoke iron-restriction in microfluidic chambers.

For microfluidic experiments, the chips were inoculated with monocultures or co-cultures adjusted to an OD_600_ of 0.5. Time-lapse imaging was performed using a Nikon Eclipse Ti2 microscope equipped with a 100× Plan Apo λ oil immersion lens, a DS-Qi2 CMOS camera, and a Perfect Focus System to prevent axial drift. Phase contrast and fluorescence images were captured every 30 min. The fluorescence of the genomically encoded E2-Crimson, GFP and mTagBFP were excited using a Sola LED illumination system with the appropriate filter cubes (E2-Crimson: λ_Ex_ = 562/40 nm, λ_Em_ = 641/84 nm; GFP: λ_Ex_ = 520/40 nm, λ_Em_ = 540/40 nm; mTagBFP: λ_Ex_ = 438/24 nm, λ_Em_ = 483/32 nm).

Initial preprocessing of the raw microscopy data was performed in Fiji (Version 1.54p). There, the images were cropped to a defined field of view of approximately 100 × 60 µm. Next, the time-lapse sequences were exported as TIFF files and uploaded to an open microscopy environment (OMERO) server.

Subsequent image analysis was implemented in custom Jupyter-based workflows adapted from previously published acia workflows [89]. Image segmentation relied on the Acia framework in conjunction with the Omnipose algorithm. Fluorescence signals enabled cell differentiation, and segmentation artifacts were excluded through a combination of intensity thresholding and size-based filtering. To further refine the results, position-based criteria were applied to omit regions corresponding to chamber boundaries. From the resulting segmentation masks, individual cell areas were quantified in regards of their observed cell area and aggregated for further analysis for each frame.

Subsequent analyses focused on the temporal progression of colony area, defined as the cumulative area of all cells per species within a chamber. Growth rates were calculated for individual time points by forming the difference quotient between the cell area present at the analyzed time point and the cell area present at the time point before.

Additionally, we characterized relative population dynamics by evaluating cell area ratios over time. The ratio of cells at a given timepoint is the fraction of the total area of an analyzed color divided by the total area of all cells present in a chamber.

### Model and simulation of microfluidic co-culture experiments

We formulated a deterministic, unstructured population model to describe the interaction between *E. coli* and *C. glutamicum* in microfluidic chambers. The system is represented by coupled ordinary differential equation (ODE) based on Monod-type kinetics. We assumed: well-mixed chamber, with no spatial gradients, no explicit glucose limitation, growth-associated metabolite exchange only, and no cell death or maintenance term. The production of enterobactin and its binding to iron is simplified into the production of an abstract product *P*_*Enterobactin*_. The set of governing reactions read:

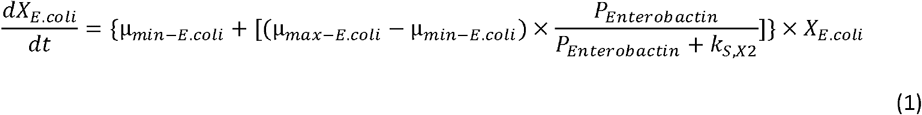

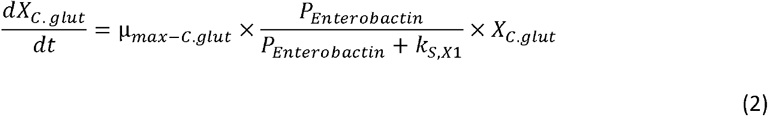

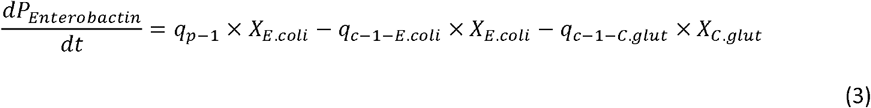

Where *μ*_*min – E*.*coli*_ is the minimal growth rate of *E. coli* in absence of *P*_*Enterobactin*_, *μ*_*max-min*_ is the maximum growth rate of *E. coli, k*_*S,X*2_ is the half saturation constant towards *P*_*Enterobactin*_ of *E. coli, k*_*S,X*1_ is the half saturation constant towards *P*_*Enterobactin*_ of *C. glutamicum, X*_*E*.*coli*_ and *X*_*C*.*glut*_ are the concentrations of both organisms, 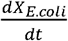 and 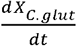 are the change of *E. coli* and *C. glutamicum* concentration and 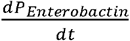 is the change of the concentration of *P*_*Enterobactin*_ In this scenario *E. coli* produces a product which itself need to take up again to stimulate growth. The production and the consumption of *P*_*Enterobactin*_ by both organisms is assumed to be growth dependent.

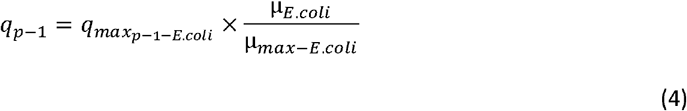

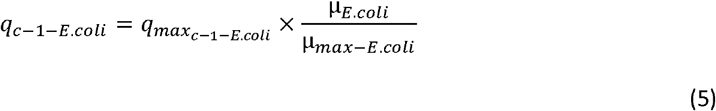

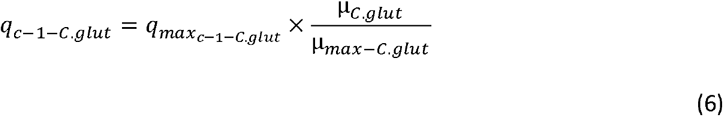

These proposed rates have been adjusted to fit qualitatively to our observed data. A complete list of all parameters is given in the supplements (Table S6 and S7).

### Parametrization

Maximum specific growth rates of both organisms were experimentally determined or obtained from literature (Table S6). The half-saturation constant for *E. coli* was taken from literature, whereas the corresponding parameter for *C. glutamicum* was varied to test the hypothesized interaction mechanism.

Production and uptake coefficients were defined relative to maximum growth rates and adjusted to reproduce qualitative experimental trends. All parameter values and units are summarized in Supplementary Table S6 and S7.

### Varying starting conditions in microfluidic experiments

To demonstrate the effects of the varying starting conditions given in large numbers of microfluidic cultivation chambers, we performed two parameter sweeps. In a first step we varied the starting concentrations of total cell dry-weight, which corresponds to different numbers of starting cells which are inside microfluidic chambers. The variation of cell dry-weight corresponds with different start cell areas of all cells in a chamber between 10 µm^2^ and 100 µm^2^.

In the second step the simulated starting cell dry weight concentration is simulated as 19 different compositions of *C. glutamicum* and *E. coli*, between 5% and 95% of the entire cell dry weight concentration. The starting concentration of *P*_*Enterobactin*_ was set, so the calculated growth rate for *E. coli* is 95% of its maximum growth rate. In total 133 individual simulations with varying starting cell areas and compositions were performed.

### Varying affinity relationships of co-culture participants

To investigate the impact of potential variants of affinity towards *P*_*Enterobactin*_ we performed another two-dimensional parameter sweep. We varied the affinity of *C. glutamicum* towards the limiting product by scaling its Monod constant *k*_*S-E-C*_ relative to the producers Monod constant *k*_*S-E-E*_ from 0.01 to 100, logarithmically scaled. The baseline assumption of 10 was explicitly modelled as well. To evaluate the system’s performance during active expansion, the mean specific growth rates for both species were calculated exclusively for the growth phase up to a total biomass of 76.15 g/L, corresponding to an observed projected cell area of 2500 µm^2^.

### Numerical integration

The model was solved numerically using Python (Version 3.8.15) and the scipy.integrate library (Version 1.9.1). The LSODA integrator was employed via the solve_ivp function. Specific growth rates were recalculated post-simulation based on the resulting concentration profiles and the governing Monod kinetics. Solutions were evaluated at 0.1 h intervals over a total simulation time of 30 h.

## Code availability

The code has been deposited in a public repository (https://jugit.fz-juelich.de/m.pesch/race-for-iron-model-code.git), ensuring full reproducibility of all results presented in this study. Additionally, the model has been deployed in a browser and can be interactively interrogated under computational-biology-aachen.github.io/mxl-web/dynamic-entrobactin.

## Acknowledgements

Our special thanks to the Institute for Biochemistry for their friendly support in laboratory equipment.

## Data availability

The datasets generated and/or analyzed during the current study are included in this published article and its supplementary information files.

## Declaration of generative AI and AI-assisted technologies in the writing

During the preparation of this work the authors used ChatGPT in order to improve language of the manuscript. After using this tool, the authors reviewed and edited the content as needed and take full responsibility for the content of the manuscript.

