## Supplementary Information for "Context-dependent siderophore exploitability shapes microbial community structure"

**Figure S1:** Full data set of plate assay on xenosiderophore utilization in *C. glutamicum.*

**Figure S2:** Spent media from *E. coli* is promoting growth of *C. glutamicum.*

**Figure S3:** Metabolomic analysis of tricarboxylic acid cycle compounds in *C. glutamicum* shows reduced citric acid production in the presence of enterobactin suggesting reduced iron limitation.

**Figure S4:** *In vitro* characterization of IronSenseR-mCherry.

**Figure S5:** *C. glutamicum* and *E. coli* duo co-cultivations demonstrate enterobactin utilization at different extents.

**Figure S6:** Duo co-cultivations with *P. putida* demonstrate pyoverdine as selfish good for *C. glutamicum* and *E. coli.*

**Figure S7:** *P. putida* is able to exploit enterobactin from *E. coli* WT and consequently tune down own siderophore production.

**Figure S8:** Growth curves and flowcytometry measurements of *E. coli*, *C. glutamicum* and *P. putida* under various iron conditions.

**Figure S9:** Representative single channel images of plate assays with the trio-consortium of *C. glutamicum*, *E. coli* and *P. putida*.

**Figure S10**: Plate assays with the trio-consortium of *C. glutamicum*, *E. coli* and *P. putida* reveals siderophore characteristics as public good and selfish good.

**Table S1:** Overview of putative siderophore-related genes in *Corynebacterium glutamicum.*

**Table S2:** Calculated ratio-dependent growth rates from microfluidic single-cell analysis growing in CGXII medium supplemented with 2% glucose and under iron limitation (3.6 µM FeSO4) based on cell area.

**Table S3:** Microbial strains used in this study.

**Table S4:** Plasmids used in this study.

**Table S5:** Primers used in this study.

**Table S6:** Description of all set parameters used for the modelling of microfluidic co-culture growth.

**Formula S1:** Derivation of conversion factor between measured cell area and volume of cells in the microfluidic system.

**Video S1:** Microfluidic cultivation of *E. coli-gfp* (yellow) and *C. glutamicum-E2-crimson* (red) (Figure 3B).

**Video S2:** Microfluidic cultivation of *E. coli ΔentC-gfp* (yellow) and *C. glutamicum-E2-crimson* (red) (Figure 3C).

**Video S3:** Microfluidic cultivation of *E. coli-gfp* (yellow), *C. glutamicum-E2-crimson* (red) and *P. putida-bfp* (blue) with high abundances of *C. glutamicum* (Figure 5A).

**Video S4:** Microfluidic cultivation of *E. coli-gfp* (yellow), *C. glutamicum-E2-crimson* (red) and *P. putida-bfp* (blue) with high abundances of *E. coli* (Figure 5A).

**Video S5:** Microfluidic cultivation of *E. coli-gfp* (yellow), *C. glutamicum-E2-crimson* (red) and *P. putida-bfp* (blue) with high abundances of *P. putida* (Figure 5A).

**Video S6:** Microfluidic cultivation of *E. coli-gfp* (yellow), *C. glutamicum-E2-crimson* (red) and *P. putida* Δ*pvdD-bfp* (blue) with high abundances of *C. glutamicum* (Figure 5B).

**Video S7:** Microfluidic cultivation of *E. coli-gfp* (yellow), *C. glutamicum-E2-crimson* (red) and *P. putida* Δ*pvdD-bfp* (blue) with high abundances of *E. coli* (Figure 5B).

**Video S8:** Microfluidic cultivation of *E. coli-gfp* (yellow), *C. glutamicum-E2-crimson* (red) and *P. putida* Δ*pvdD-bfp* (blue) with high abundances of *P. putida* (Figure 5B).

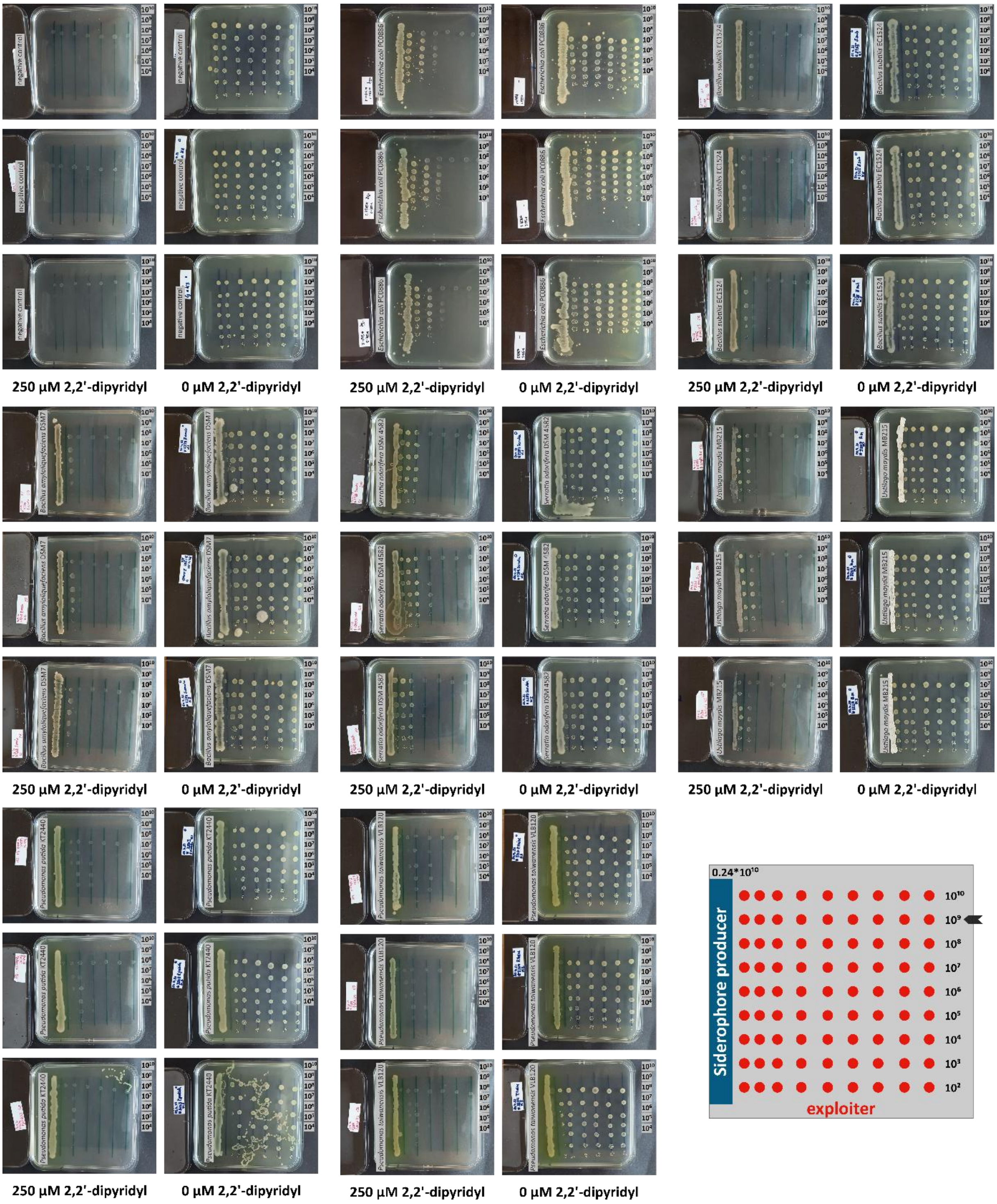


**Figure S1: Full data set of plate assay on xenosiderophore utilization in *C. glutamicum.*** Growth promotion of iron-starved *C. glutamicum* by seven known siderophore-producing strains was tested on double-layer LB agar containing 250 µM 2,2′-dipyridyl (DIP) as an iron chelator. Plates without chelator served as control (0 µM 2,2´-dipyridyl). *C. glutamicum* cell suspensions with different cell densities were spotted (3 µL, $10^4$-$10^9$ cells/mL) at different distances to the pre-grown lane of siderophore producer (40 µL, $0.24*10^{10}$ cells/mL) and incubated at 30 °C for two days. The experiment was performed in independent biological duplicates or triplicates.

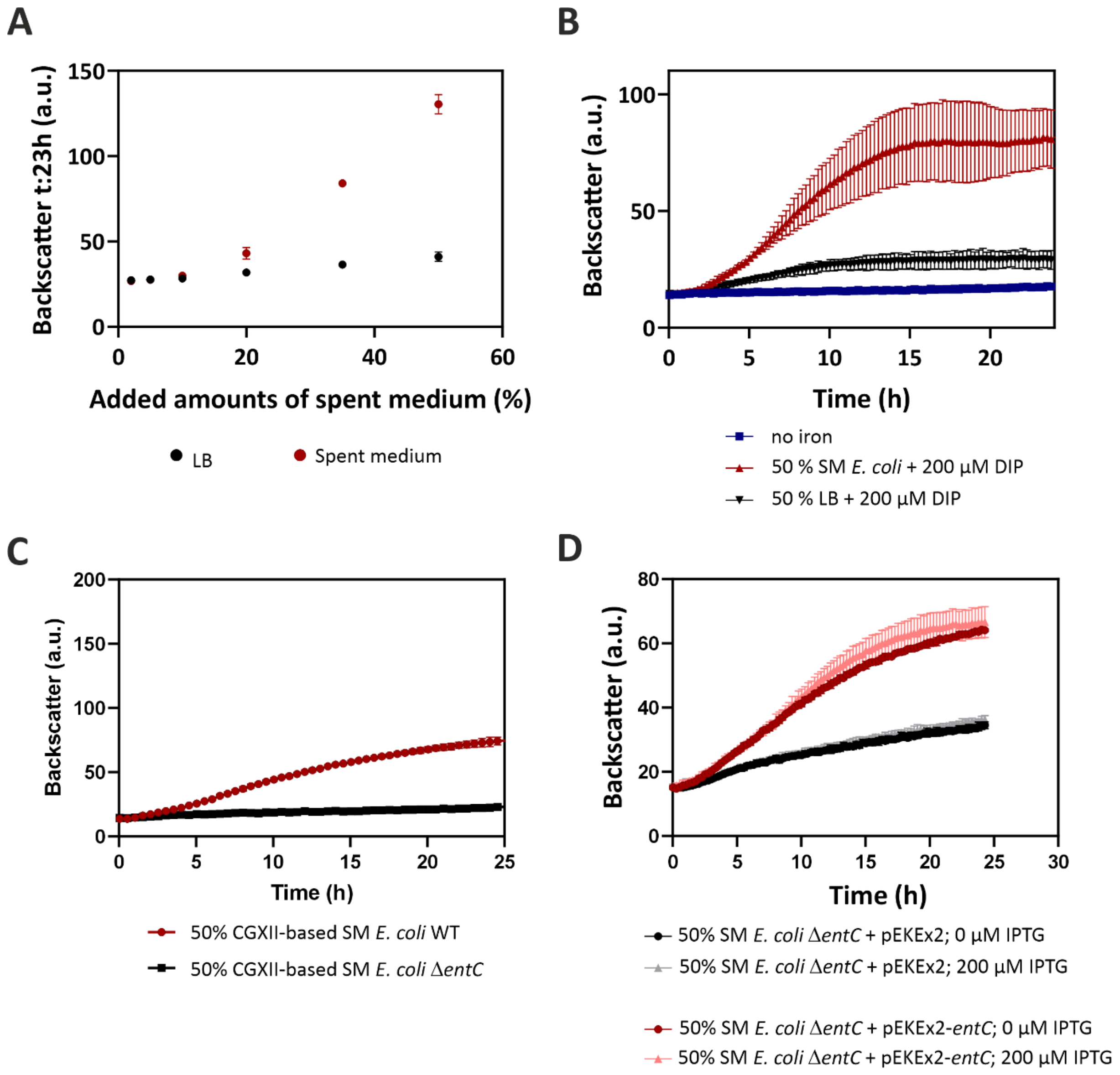


**Figure S2: Spent medium from *E. coli* is promoting growth of *C. glutamicum*.** Further control experiments underlining the growth-promoting effects of the spent medium (SM) from *E. coli*. **(A)** Dose-response curve at time point 23 h corresponding to growth of iron-starved *C. glutamicum* in the presence of varying amounts of LB or spent medium from *E. coli*. **(B)** Growth of *C. glutamicum* growing in the presence of 50% 2-fold CGXII supplemented with 2% glucose and without iron. Further 50% were either SM supplemented with 200 µM 2,2'-dipyridyl (DIP) (light red) or without (dark red). As control, 50% LB with 200 µM DIP (black) and also 50% CGXII without any iron (blue) were supplemented. **(C)** Comparing growth of *C. glutamicum* on SM from *E. coli* WT and the siderophore-deficient Δ*entC* variant using CGXII-based SM, instead of LB-based (Figure 2B, main text). **(D)** *E. coli* Δ*entC* strain was transformed with either empty plasmid (pEKEx2) or a plasmid harboring *entC* under the IPTG-inducible $P_{tac}$ promoter (pEKEx2-*entC*). SM from both strains was harvested. *C. glutamicum* was cultivated in 50% 2-fold CGXII supplemented with 2% glucose and no iron, but 25 µg/mL kanamycin and either 0 or 200 µM IPTG, as well as 50% LB-SM of these respective *E. coli* strains including the *entC*-deficient (shades of black) or *entC*-overexpression (shades of red) strains. Backscatter (a.u.) values were measured in a microtiter cultivation system with *n* = 3 biological replicates.

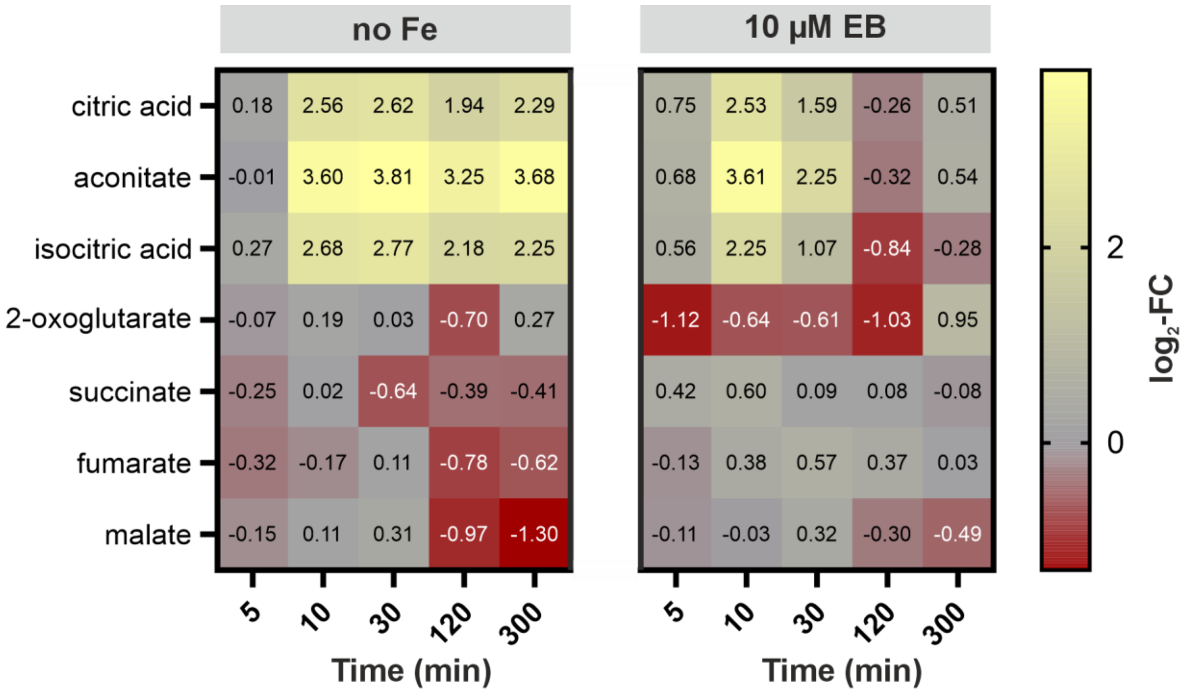


**Figure S3: Metabolomic analysis of tricarboxylic acid cycle compounds in *C. glutamicum* shows reduced citric acid production in the presence of enterobactin suggesting reduced iron limitation.** The heatmap represents the $log_2$-fold change (FC) of the normalized area (a.u.) of the MS-measurements of the no Fe or 10 µM enterobactin condition in relation to the full iron (36 µM $FeSO_4$) condition. *C. glutamicum* was cultivated in CGXII supplemented with 2% glucose and either (i) no added $FeSO_4$ but 150 µM DIP (no Fe), or (ii) 3.6 µM $FeSO_4$, 150 µM DIP and 10 µM enterobactin (10 µM EB) or (iii) 36 µM $FeSO_4$ (full iron) and no DIP (iron saturated condition). *n*= 4 biological replicates.

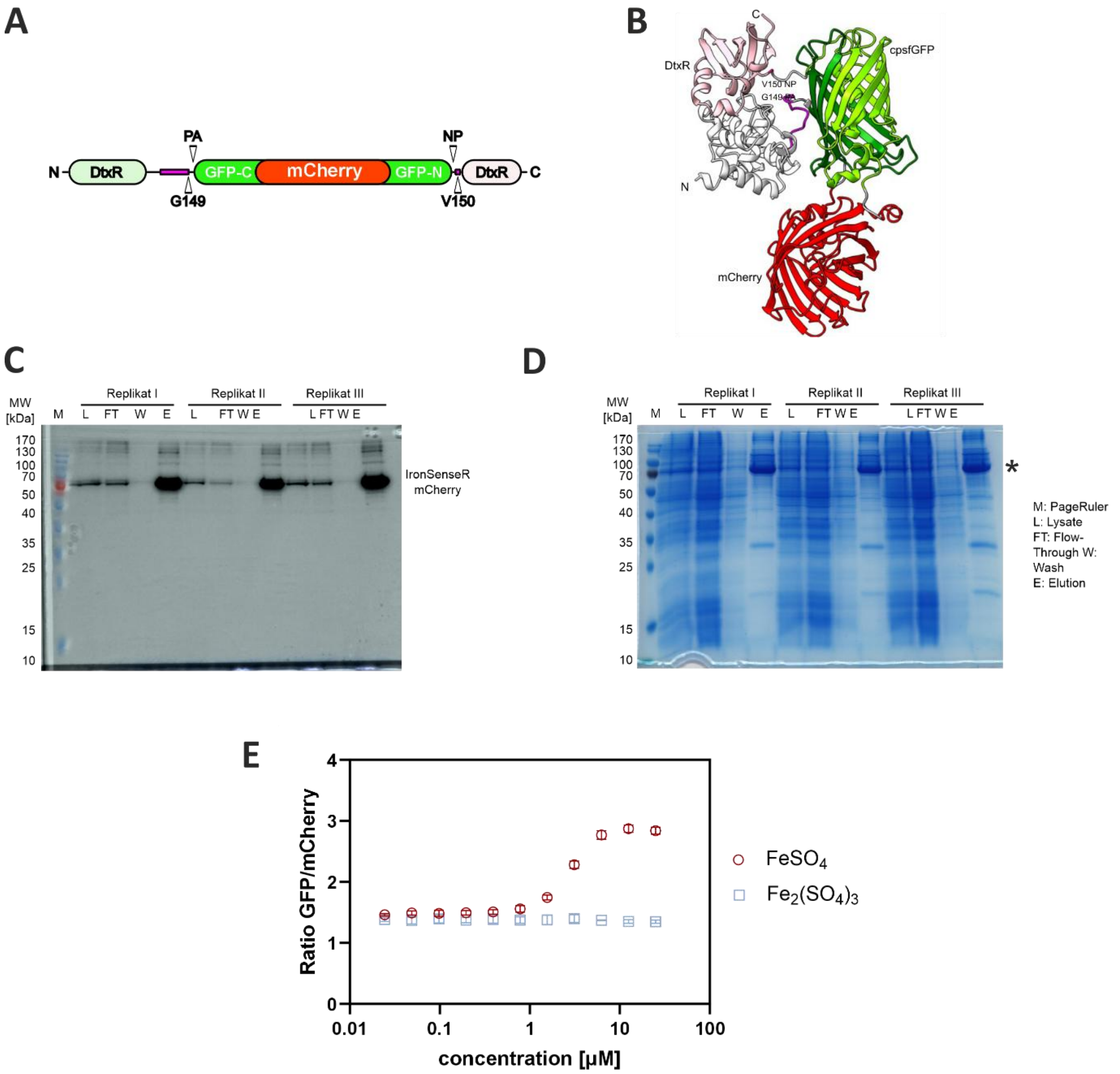


**Figure S4: *In vitro* characterization of IronSenseR-mCherry. (A)** Molecular architecture and **(B)** AlphaFold3 model of Matryoshka biosensor IronSenseR-mCherry for *in vivo* $Fe^{2+}$-detection in *C. glutamicum* and *P. putida*. According to Papadopoulos, Anlauf et al. (2026) the fluorescence intensity of the cpsfGFP domain increases with increasing $Fe^{2+}$-concentrations. The biosensor expression was conducted in the bacterial expression host *E. coli* and protein fractions were analyzed by SDS-PAGE after purification via affinity chromatography. **(C)** Coomassie stained images of the SDS-PAGE and **(D)** in-gel fluorescence of the SDS-PAGE of Matryoshka biosensor, IronSenseR-mCherry. The prominent band at approximately 80 kDa labeled with asterisk (*) indicates the biosensor. L: Cell lysate, FT: Flow through, W: Wash fraction, E: Elution fraction from affinity chromatography. The molecular weight is indicated in kDa. **(E)** The specificity of IronSenseR-mCherry for ferrous iron ($FeSO_4$) and ferric iron ($Fe_2(SO_4)_3$) was analyzed *in vitro*. Concentration of metal ions utilized for titration ranging from 0-25 µM. The experiments conducted in *n* = 3 biological replicates, mean values and the standard deviation (SD) is indicated. Calculated $K_D$-value derived from curve fit to $FeSO_4$ is 2.369 ± 0.143 µM.

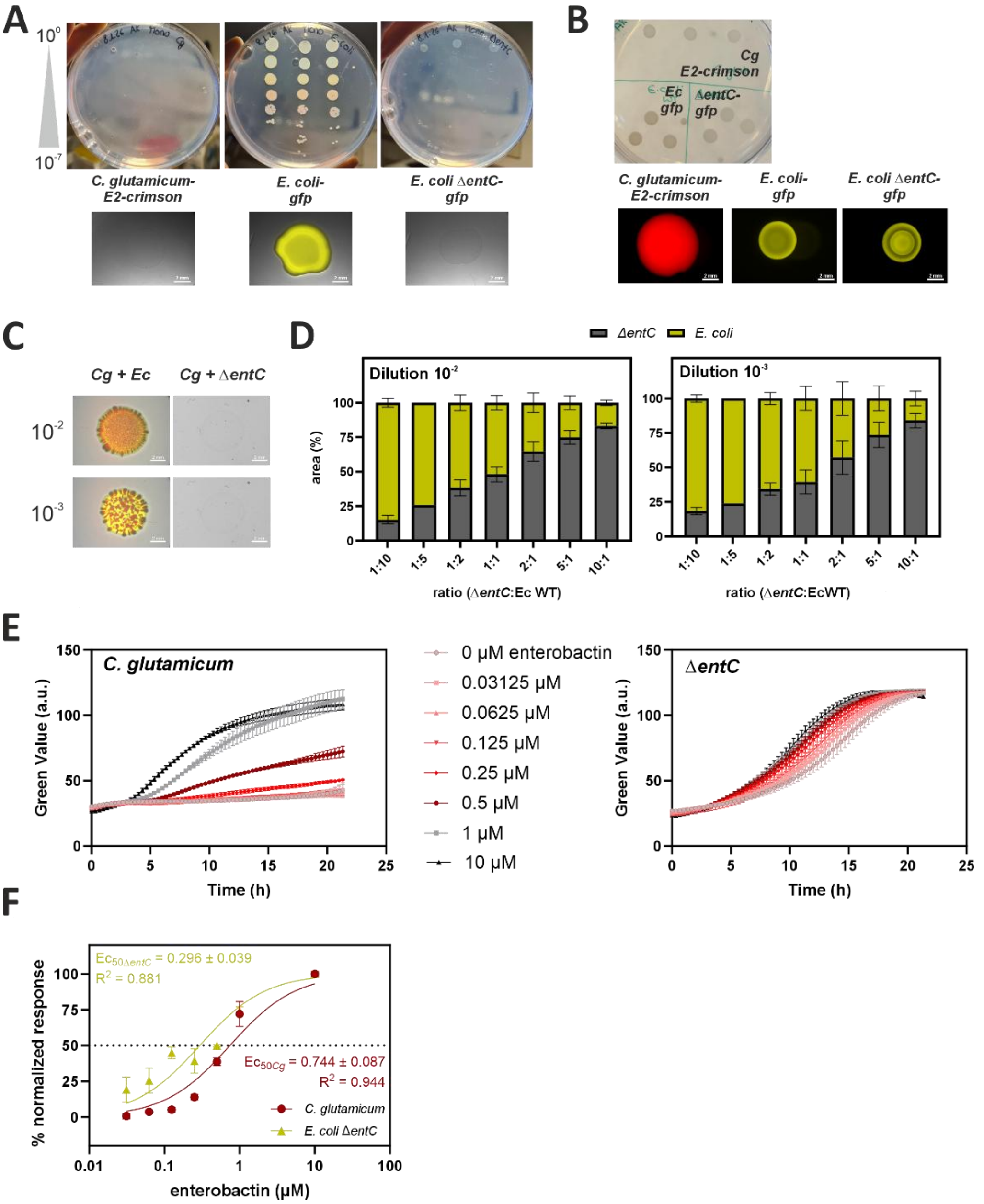


**Figure S5: *C. glutamicum* and *E. coli* duo co-cultivations demonstrate enterobactin utilization at different extents. (A)** Cultivation of *C. glutamicum*-E2-*crimson*, *E. coli-gfp* and *E. coli ΔentC-gfp* on CGXII agar plates containing 2% glucose, 3.6 µM $FeSO_4$ and 100 µM DIP in different dilutions in $n$ = 3 biological replicates (top) and single drop analysis using a stereomicroscope (bottom). **(B)** Same strains in the presence of 36 µM $FeSO_4$ and no DIP, representing a sufficient iron supply. **(C)** Mixed drop (1:1) co-cultivation of *C. glutamicum* (Cg) with *E. coli* WT (Ec) or *E. coli ΔentC* on CGXII plates with 2% glucose, 3.6 µM $FeSO_4$ and 100 µM DIP. **(D)** Quantified area in percentage for co-cultivation of *E. coli* WT-*gfp* (yellow) and *E. coli ΔentC* (grey) in varying ratios as depicted on the x-axis at two different dilutions ($10^{-2}$ and $10^{-3}$). $n$ = 3 biological replicates. **(E)** Dose-response curves of *C. glutamicum* (left) and *E. coli ΔentC* (right) growing in CGXII supplemented with 1% glucose and 3.6 µM $FeSO_4$ as well as respective, increasing amounts of enterobactin. Growth was monitored in a shaker-incubator Growth Profiler (EnzyScreen, Netherlands). **(F)** $Ec_{50}$ value derived from normalized growth rates (%) and a non-linear curve fit via GraphPad Prism version 10.5.0 (GraphPad Software, USA).

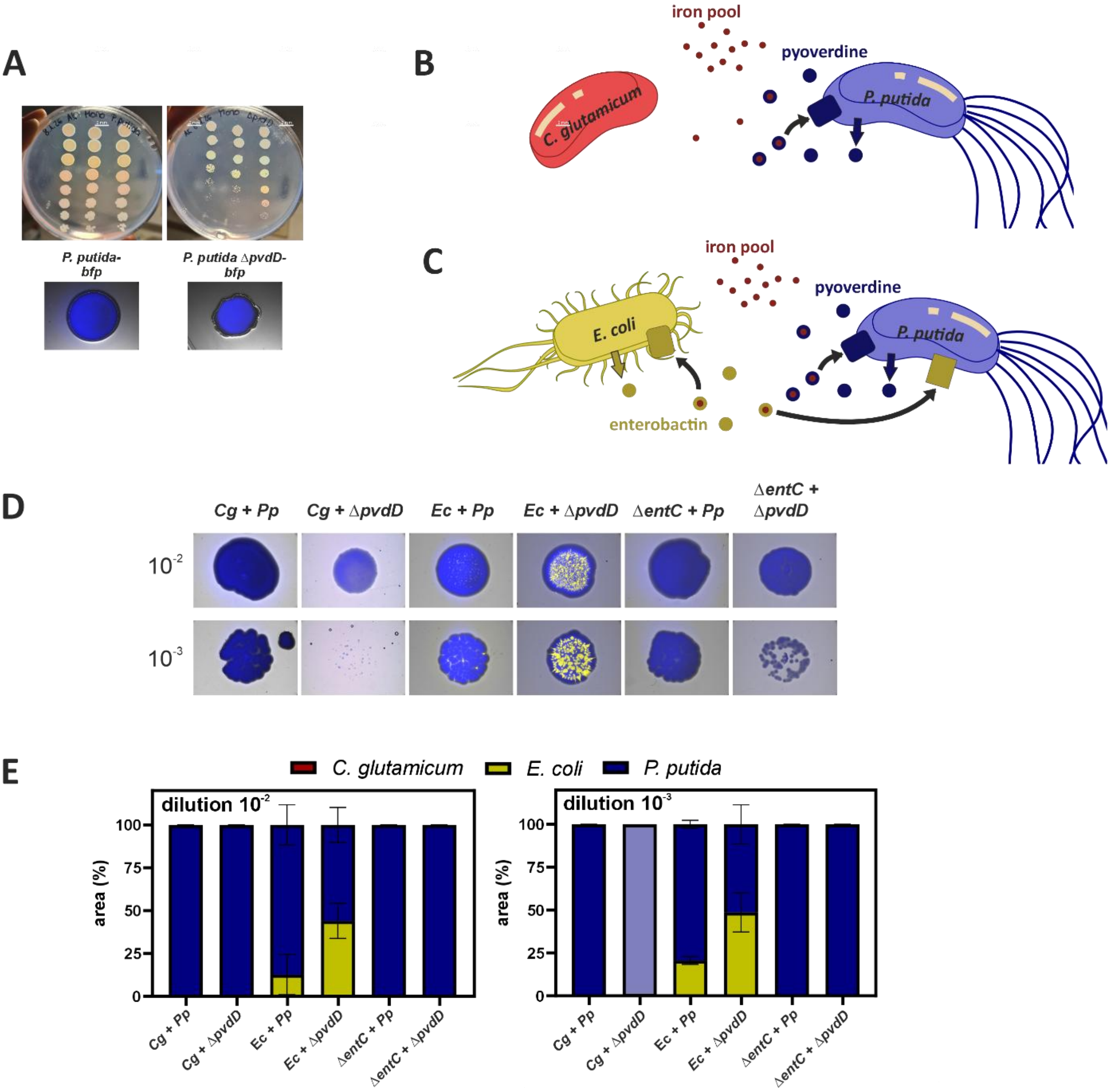


**Figure S6: Duo co-cultivations with *P. putida* demonstrate pyoverdine as selfish good for *C. glutamicum* and *E. coli*. (A)** Cultivation of *P. putida-bfp* and *P. putida ΔpvdD-bfp* on CGXII agar plates containing 2% glucose, 3.6 µM $FeSO_4$ and 100 µM DIP in increasing 10-fold dilutions in *n* = 3 biological replicates (top) and single drop analysis using a stereomicroscope (bottom). **(B)** Schematic depiction of siderophore-mediated interaction between *C. glutamicum* (non-producer) and *P. putida* (pyoverdine producer). **(C)** Schematic depiction of siderophore-mediated interaction between *E. coli* (enterobactin producer) and *P. putida* (pyoverdine producer and enterobactin exploiter, also see Figure S9). **(D)** Mixed drop (1:1) duo co-cultivation of *P. putida-bfp* (Pp) or *P. putida ΔpvdD-bfp* in all combinations with *C. glutamicum*-E2-*crimson* (Cg), *E. coli* WT-*gfp* (Ec) or *E. coli ΔentC-gfp* on CGXII plates with 2% glucose, 3.6 µM $FeSO_4$ and 100 µM DIP in two different dilutions ($10^{-2}$ and $10^{-3}$). **(E)** Quantified area in percentage for co-cultivation as shown representatively in (D). Note: Cg + Δ*pvdD* of $10^{-3}$ is depicted in light blue, because nearly no growth was observed (compare (D)). *n* = 3 biological replicates.

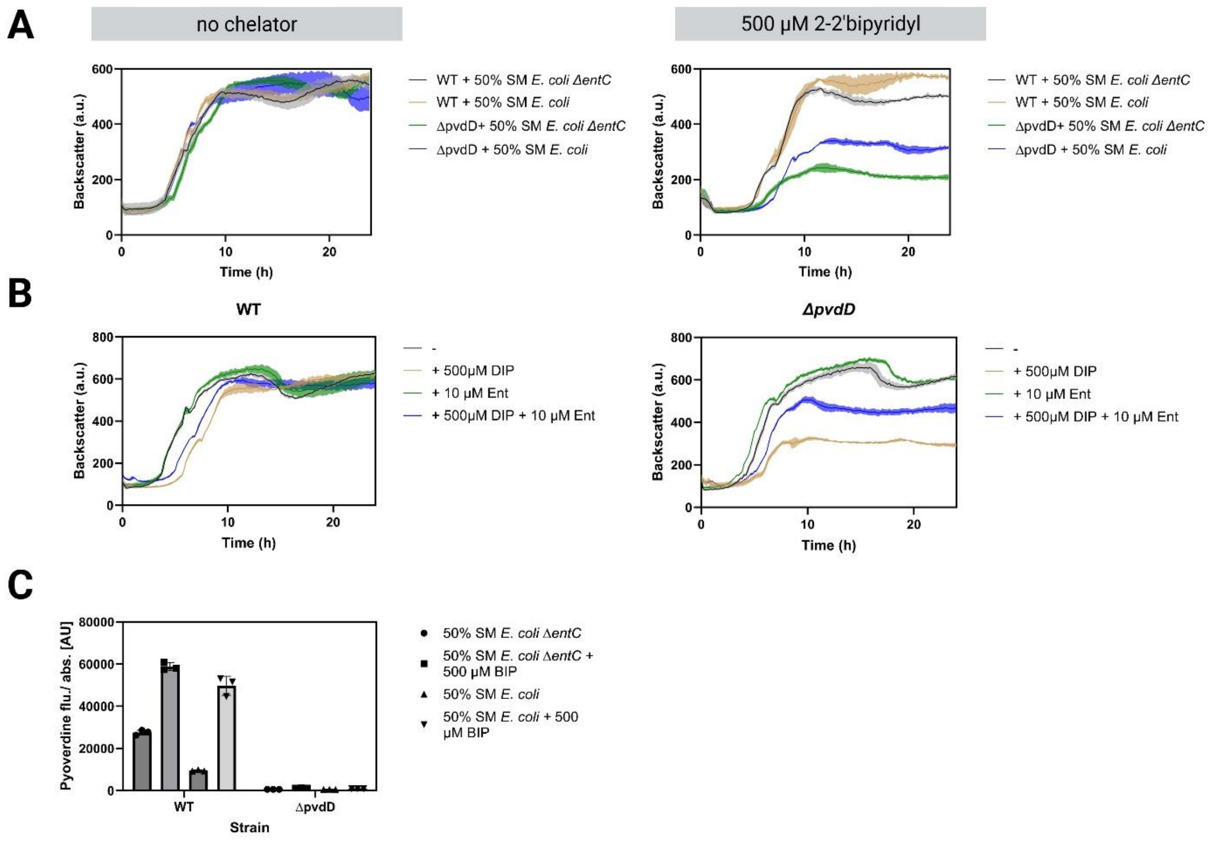


**Figure S7: *P. putida* is able to exploit enterobactin from *E. coli* WT and consequently tune down own siderophore production. (A)** Growth of *P. putida* WT (grey & beige) and *P. putida ΔpvdD* (green & blue) in LB with the addition of 50% *E. coli* spent medium (either from WT or Δ*entC*), with or without the addition of 500 µM DIP. **(B)** Growth of *P. putida* WT (left) and *P. putida ΔpvdD* (right) in LB with the addition of either 500 µM DIP (beige), 10 µM enterobactin (green) or both (blue). As control, LB without any additions was used (grey). **(C)** Pyoverdine fluorescence ($\lambda_{ex}$ = 398 nm, $\lambda_{em}$ = 455 nm) of cultures after cultivation with *E. coli* spent media.

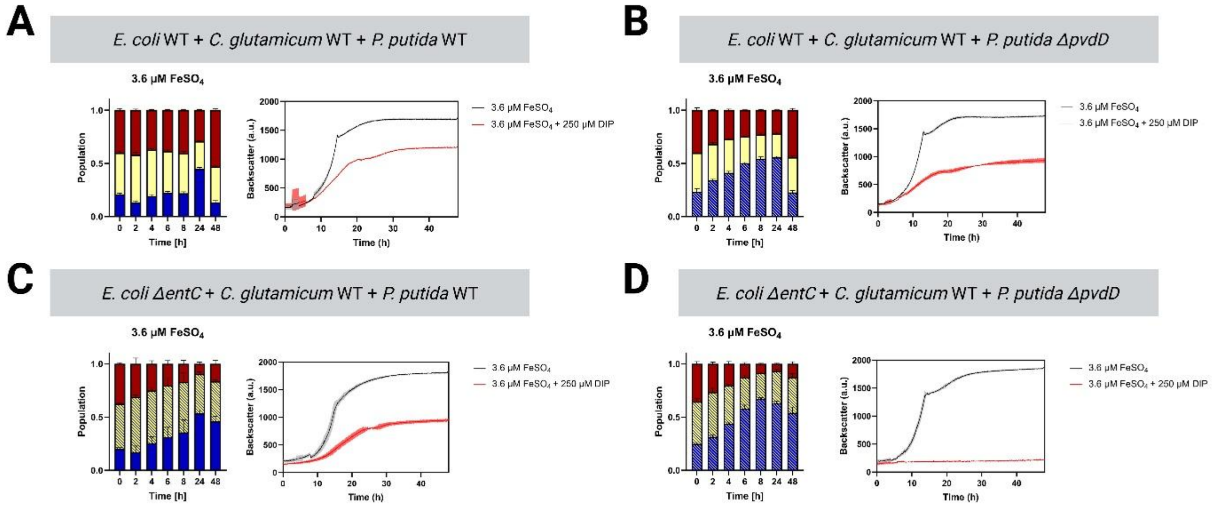


**Figure S8: Growth curves and flow cytometry measurements of *E. coli*, *C. glutamicum* and *P. putida* under various iron conditions. (A-D)** Bar charts showing populations of trio cultures grown in CGXII (2% glucose + 3.6 µM $FeSO_4$) measured via flow cytometry on the left and growth curves of the same cultures as well as the + DIP control on the right.

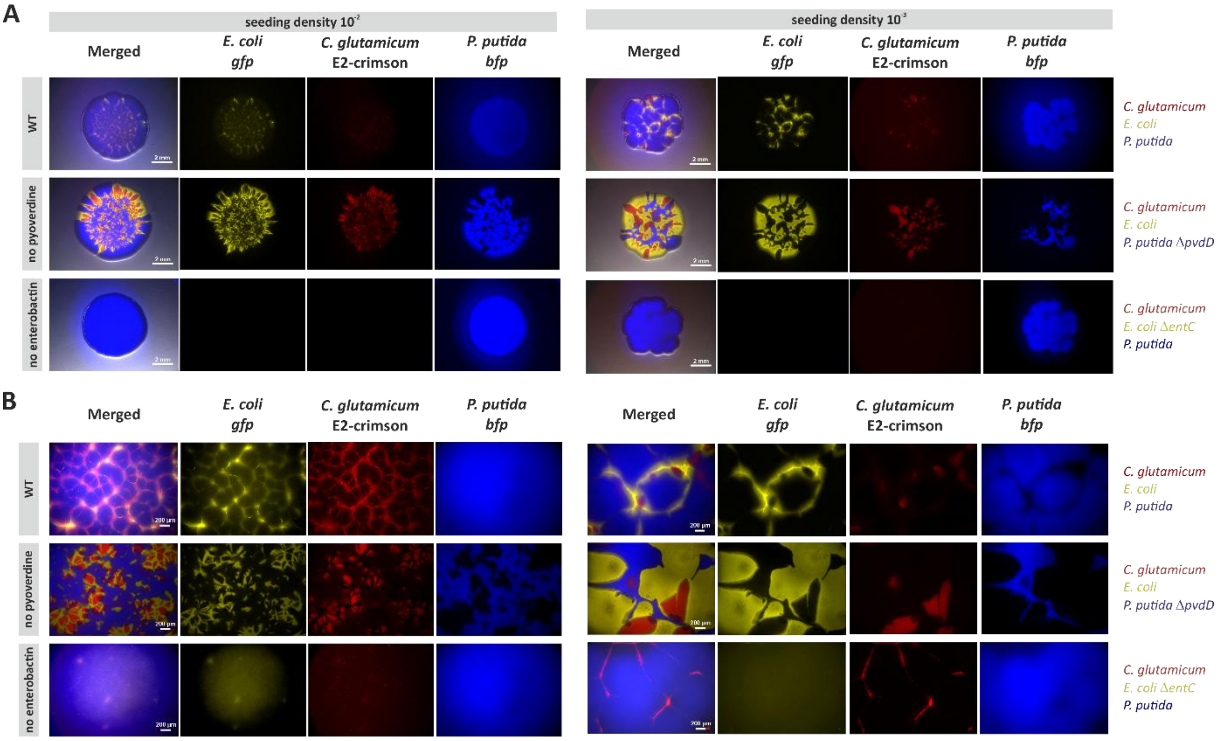


**Figure S9: Representative single channel images of plate assays with the trio-consortium of *C. glutamicum*, *E. coli* and *P. putida*.** Single channel images for the representative stereomicroscopy pictures of mixed drop assay on iron-restricted plates as shown in Figure 4G with *C. glutamicum-E2-crimson* (red), *E. coli-gfp* (yellow) and *P. putida-bfp* (blue), as well as siderophore-deficient mutants, as indicated in two different seeding densities in a 1:1:1 ratio and **(A)** 16x vs. **(B)** 80x zoom.

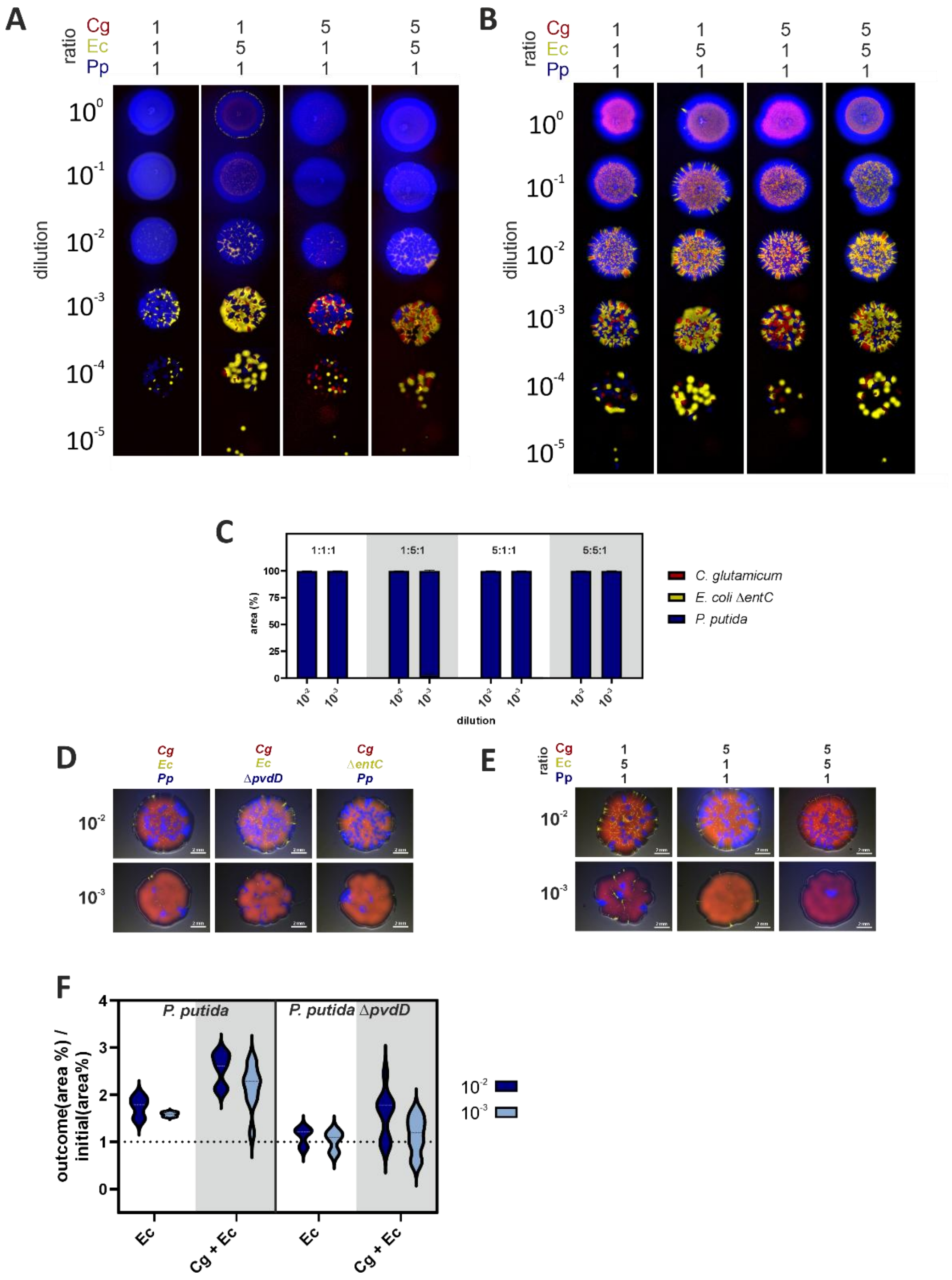


**Figure S10: Plate assays with the trio-consortium of *C. glutamicum*, *E. coli* and *P. putida* reveals siderophore characteristics as public good and selfish good. (A)** Mixed drop trio co-cultivation of *P. putida-bfp* (Pp) or **(B)** *P. putida* Δ*pvdD-bfp* with *C. glutamicum*-E2-*crimson* (Cg) and *E. coli* WT-*gfp* (Ec), in different ratios as depicted on top, on CGXII plates with 2% glucose, 3.6 µM $FeSO_4$ and 100 µM DIP in different dilutions ($10^0$ to $10^{-5}$). **(C)** Quantified area in percentage for no-enterobactin trio co-cultivation of *C. glutamicum*, *E. coli* Δ*entC-gfp* and *P. putida* at varying ratios in two different dilutions ($10^{-2}$ and $10^{-3}$) on iron-limited CGXII agar plates. *n* = 3 biological replicates. **(D)** Trio-cultivations of Cg-Ec-Pp and the siderophore-deficient variants on CGXII agar plates with 2% glucose, 36 µM $FeSO_4$ and PCA as siderophore-independent iron excess control. **(E)** WT variants in different ratios at iron excess control. One representative of *n* = 3 biological replicates. **(F)** *P. putida* WT and Δ*pvdD* outcome analysis with a ratio of *P. putida* outcome over initial area in percentage for two seeding densities ($10^{-2}$ and $10^{-3}$) in duo and trio co-cultivations.

**Table S1: Overview of putative siderophore-related genes in *Corynebacterium glutamicum.***

| Locus | Start | Stop | Annotation | up-/down-regulated in response to iron limitation* |
|---|---|---|---|---|
| NCgl0036 | 37245 | 38201 | putative iron-siderophore ABC transporter, permease protein | down |
| NCgl0037 | 38202 | 38981 | putative iron-siderophore ABC transporter, ATP-binding protein | |
| NCgl0038 | 38978 | 39802 | putative iron-chelator utilization protein | |
| NCgl0329 | 352693 | 353640 | putative ironIII dicitrate ABC transporter, secreted siderophore-binding lipoprotein | up |
| NCgl0411 | 448127 | 449179 | putative spermidine/putrescine/ironIII ABC transporter, ATPase subunit | down |
| NCgl0412 | 449180 | 450826 | putative spermidine/putrescine/ironIII ABC transporter, permease subunit | |
| NCgl0413 | 450834 | 451913 | putative spermidine/putrescine/ironIII ABC transporter, substrate-binding lipoprotein | |
| NCgl0482 | 528776 | 529570 | putative siderophore ABC transporter, ATP-binding protein | up |
| NCgl0483 | 529589 | 530755 | putative siderophore ABC transporter, permease protein | |
| NCgl0484 | 530745 | 531791 | putative siderophore ABC transporter, permease protein | |
| NCgl0618 | 659543 | 660541 | putative iron-siderophore ABC transporter, substrate-binding lipoprotein | up |
| NCgl0635 | 680128 | 681027 | putative cytoplasmic siderophore-interacting protein | up |
| NCgl0636 | 681037 | 681846 | putative iron-siderophore ABC transporter, ATP-binding protein | |
| NCgl0637 | 681843 | 682871 | putative iron-siderophore ABC transporter, permease subunit | |
| NCgl0638 | 682868 | 683863 | putative iron-siderophore ABC transporter, permease subunit | |
| NCgl0639 | 683873 | 684925 | putative iron-siderophore ABC transporter, secreted siderophore-binding lipoprotein | |
| NCgl0644 | 688913 | 689890 | putative iron-siderophore ABC transporter, substrate-binding lipoprotein | n.d. |
| NCgl0645 | 689914 | 690696 | putative iron-siderophore ABC transporter, ATPase subunit | |
| NCgl0646 | 690703 | 691722 | putative iron-siderophore ABC transporter, permease subunit | |
| NCgl0773 | 848496 | 849323 | putative cytoplasmic siderophore-interacting protein | up |
| NCgl0774 | 849323 | 850243 | putative secreted siderophore-binding lipoprotein | |

| | | | | |
|---|---|---|---|---|
| NCgl0776 | 851351 | 852367 | putative iron-siderophore ABC transporter, substrate-binding lipoprotein | up |
| NCgl0777 | 52618 | 853619 | putative iron-siderophore ABC transporter, permease subunit | up |
| NCgl0778 | 853612 | 854727 | putative iron-siderophore ABC transporter, permease subunit | |
| NCgl0779 | 854724 | 855479 | putative iron-siderophore ABC transporter, ATPase subunit | |
| NCgl1200 | 1313270 | 1314121 | putative cytoplasmic siderophore-interacting protein | up |
| NCgl1209 | 1322393 | 1323409 | putative secreted siderophore-binding lipoprotein | up |
| NCgl1395 | 1529486 | 1530223 | putative cytoplasmic siderophore-interacting protein | n.d. |
| NCgl1564 | 1721780 | 1722856 | putative iron-siderophore ABC transporter, substrate-binding lipoprotein | n.d. |
| NCgl1565 | 1722870 | 1723829 | putative iron-siderophore ABC transporter, ATPase subunit | |
| NCgl1566 | 1723826 | 1724581 | putative iron-siderophore ABC transporter, permease subunit | |
| NCgl1905 | 2089215 | 2089868 | phosphopantetheinyl transferase, iron-chelating complex subunit | n.d. |
| NCgl1959 | 2148225 | 2149169 | putative ironIII dicitrate ABC transporter, substrate-binding lipoprotein | up |
| NCgl2031 | 2229096 | 2229896 | putative ironIII dicitrate ABC transporter, ATPase subunit | n.d. |
| NCgl2032 | 2229897 | 2230937 | putative ironIII dicitrate ABC transporter, permease subunit | |
| NCgl2033 | 2230944 | 2231933 | putative ironIII dicitrate ABC transporter, substrate-binding lipoprotein | |
| NCgl2970 | 3284397 | 3285425 | Putative ironIII dicitrate ABC transporter, substrate-binding lipoprotein | up |

*(Küberl, Mengus-Kaya et al. 2020), comparison of low (1 µM) and high (36 µM) $FeSO_4$ levels, genes with a ≥ 2-fold altered mRNA ratio and a ρ value of ≤ 0.05.

**Table S2: Calculated ratio-dependent growth rates from microfluidic single-cell analysis growing in CGXII medium supplemented with 1% glucose and under iron limitation (3.6 µM $FeSO_4$) based on cell area.**

| Culture type | Organism | Partner 1 | Partner 2 | µ ± SD ($h^{-1}$) | *N* (chambers) |
|---|---|---|---|---|---|
| Mono | *C. glutamicum* WT | – | – | 0.01 ± 0.01 | 26 |
| Mono | *E. coli* WT | – | – | 0.26 ± 0.03 | 43 |
| Mono | *E. coli* Δ*entC* | – | – | 0.21 ± 0.02 | 24 |
| Mono | *P. putida* WT | – | – | 0.35 ± 0.02 | 12 |
| Mono | *P. putida* Δ*pvdD* | – | – | 0.27 ± 0.01 | 10 |
| Co | *C. glutamicum* WT | *E. coli* WT | – | 0.29 ± 0.03 | 24 |
| Co | *E. coli* WT | *C. glutamicum* WT | – | 0.23 ± 0.03 | 24 |
| Co | *C. glutamicum* WT | *E. coli* Δ*entC* | – | 0.03 ± 0.01 | 27 |
| Co | *E. coli* Δ*entC* | *C. glutamicum* WT | – | 0.17 ± 0.02 | 27 |
| Co | *E. coli* WT | *P. putida* WT | - | 0.14 ± 0.08 | 34 |
| Co | *P. putida* WT | *E. coli* WT | - | 0.32 ± 0.03 | 34 |
| Co | *E. coli* WT | *P. putida* Δ*pvdD* | - | 0.20 ± 0.08 | 32 |
| Co | *P. putida* Δ*pvdD* | *E. coli* WT | - | 0.18 ± 0.10 | 32 |
| Co | *E. coli* Δ*entC* | *P. putida* WT | - | 0.00 ± 0.13 | 22 |
| Co | *P. putida* WT | *E. coli* Δ*entC* | - | 0.33 ± 0.03 | 22 |
| Co | *E. coli* Δ*entC* | *P. putida* Δ*pvdD* | - | 0.05 ± 0.05 | 42 |
| Co | *P. putida* Δ*pvdD* | *E. coli* Δ*entC* | - | 0.14 ± 0.07 | 42 |
| Tri | *C. glutamicum* WT | *E. coli* WT | *P. putida* WT | 0.14 ± 0.08 | 36 |
| Tri | *E. coli* WT | *C. glutamicum* WT | *P. putida* WT | 0.20 ± 0.10 | 36 |
| Tri | *P. putida* WT | *E. coli* WT | *C. glutamicum* WT | 0.26 ± 0.12 | 36 |
| Tri | *C. glutamicum* WT | *E. coli* WT | *P. putida* Δ*pvdD* | 0.16 ± 0.09 | 26 |
| Tri | *E. coli* WT | *C. glutamicum* WT | *P. putida* Δ*pvdD* | 0.25 ± 0.02 | 26 |
| Tri | *P. putida* Δ*pvdD* | *E. coli* WT | *C. glutamicum* WT | 0.22 ± 0.07 | 26 |

**Table S3: Microbial strains used in this study.**

| Strain | Relevant characteristics | Reference | Purpose |
|---|---|---|---|
| ***Corynebacterium glutamicum*** | | | |
| ATCC 13032 | Biotin-auxotrophic wild type | DSMZ culture collection: DSM20300 (Kinoshita, Udaka et al. 2004) | *C. glutamicum* WT, siderophore exploiter |
| WT::cg1121/22_$P_{tet}$ *crimson* | ATCC 13032 derivative with an inserted *e2-crimson* gene under control of the constitutive $P_{tet}$ promoter (-*tetR*, constitutive), integrated in the cg1121/22 intergenic region | This study | *C. glutamicum* WT, E2-crimson-labeled |
| ***Escherichia coli*** | | | |
| DH5α | F⁻⁻ *supE44* Δ*lacU169* (ϕ80*lacZ*DM15) *hsdR17 recA1 endA1 gyrA96 thi-1 relA1* | Invitrogen | Cloning procedures |
| *E. coli* Pir2 | F - Δlac169 rpoS (Am) robA1 creC510 hsdR514 endA reacA1 uidA (ΔMlui)::pir | Life Technologies | Cloning of plasmids with R6K ori |
| *E. coli* HB101 | F − mcrB mrr hsdS20(rB− mB− ) recA13 leuB6 ara-14 proA2 lacY1 galK2 xyl-5 mtl-1 rpsL20(SmR ) gln V44λ− | Boyer and Roulland-Dussoix (1969) | Conjugation |
| PC 0886 | lambda⁻ F⁻ *sup*⁻ | DSMZ culture collection: DSM 13127 | *E. coli* WT, siderophore producer |
| PC 0886 Δ*entC* | PC 0886 derivative with in-frame deletion of the *entC* gene lambda⁻ F⁻ *sup*⁻ Δ*entC* | This study | Non-siderophore producer |
| PC 0886 *nupG*::*neoR*_ oopT_$P_{J23150}$ -*gfpmut1* | PC 0886 derivative with an inserted *neoR* and *gfpmut1* gene under control of $P_{J23150}$, separated by oopT and integrated behind the *nupG* locus | This study | *E. coli* WT, siderophore producer, Gfpmut1-labeled |
| PC 0886 Δ*entC* *nupG*::*neoR*_ oopT_$P_{J23150}$ -*gfpmut1* | PC 0886 Δ*entC* derivative with an inserted *neoR* and *gfpmut1* gene under control of $P_{J23150}$, separated by oopT and integrated behind the *nupG* locus | This study | *E. coli* Δ*entC*, non-siderophore producer, Gfpmut1-labeled |
| ***Pseudomonas putida*** | | | |
| KT2440 | Wild-type *P. putida*, Pyoverdine producer | ATCC culture collection: ATCC 47054 (Nelson, Weinel et al. 2002) | Siderophore producer |
| Δ*pvdD* | *P. putida* KT2440 with in-frame deletion of *pvdD* gene | Paik et al. unpublished | Non-siderophore producer |

| | | | |
|---|---|---|---|
| KT2440-bfp | *P. putida KT2440* with genomic attTn7 integration of: $P_{em7}$ *mTagBFP, GmR* | Paik et al. unpublished | *P. puitda* WT, siderophore producer, mTagBFP-labeled |
| Δ*pvdD-bfp* | *P. putida* KT2440 Δ*pvdD* with genomic attTn7 integration of: $P_{em7}$ *mTagBFP, GmR* | Paik et al. unpublished | *P. puitda* KT2440 Δ*pvdD* siderophore producer, mTagBFP-labeled |
| **Other microorganisms** | | | |
| *Serratia odorifera* ATCC 33077 | Wild-type *S. odorifera*, catechol-type siderophore producer | DSMZ culture collection: DSM 4582 | Siderophore producer |
| *Bacillus subtilis* EC1524 | Wild-type *B. subtilis*, catechol-type siderophore producer | John Innes Centre, Norwich, Great Britain | Siderophore producer |
| *Bacillus amyloliquefaciens* ATCC 23350 | Wild-type *B. amyloliquefaciens*, catechol-type siderophore producer | DSMZ culture collection: DSM7 | Siderophore producer |
| *Pseudomonas taiwanensis* VLB120 | Wild-type *P. taiwanensis*, mixes-type siderophore producer; styrene prototroph | (Panke, Witholt et al. 1998) | Siderophore producer |
| *Ustilago maydis* MB215 | Wild-type *U. maydis*, hydroxamate-type siderophore producer | DSMZ culture collection: DSM17144 | Siderophore producer |

**Table S4: Plasmids used in this study.**

| Plasmid | Relevant characteristics | Reference |
|---|---|---|
| pJC1 | $Kan^R$, $Amp^R$; $oriV_{C.g.}$, $oriV_{E.c.}$<br>*C. glutamicum*/*E. coli* shuttle vector | (Cremer, Eggeling et al. 1990) |
| pJC1-*venus*-term | $Kan^R$, $oriV_{C.g.}$, $oriV_{E.c.}$<br>pJC1 derivative carrying the *venus* coding sequence, followed by a *Bacillus subtilis* terminator sequence | (Baumgart, Luder et al. 2013) |
| pJC1-Δ*entC*_disruption-cassette | $Kan^R$, $oriV_{C.g.}$, $oriV_{E.c.}$<br>pJC1 derivative carrying the disruption cassette for in-frame deletion of *entC*, consisting of approximately 200 nt homologous up- and downstream regions flanking the *neoR* gene | This work |
| pKD13 | $Amp^R$, $Kan^R$, $oriR_{6K\gamma}$, *tL3LAM*, *rgnB*, *neoR*<br>Template plasmid for the amplification of *neoR* gene flanked by the frt sites (disruption cassette) | (Datsenko and Wanner 2000) |
| pKD46 | $Amp^R$, $oriR_{101}$, *repA101ts*, *araC, bet*, *gam*, *exo*<br>Temperature sensitive plasmid for L-arabinoase-inducible expression of the lambda red recombinase system | (Datsenko and Wanner 2000) |
| pCP20 | $Amp^R$, $Cat^R$, *ts-rep*, *cl857*, *flp*<br>Temperature sensitive plasmid for constitutive expression of FLP recombinase | (Cherepanov and Wackernagel 1995) |
| pJC1-*nupG* LF-BcuI-*nupG* RF_1S | $Kan^R$, $oriV_{C.g.}$, $oriV_{E.c.}$<br>pJC1 derivative carrying the left and right homologous regions (200-230 nt) of the integration site downstream of *nupG*, separated by a BcuI site.<br>Helper plasmid for reporter integration in *E. coli* no. 1. | This work |
| pJC1-*nupG* LF-*neoR*-oopT-BcuI-*nupG* RF_2S | $Kan^R$, $oriV_{C.g.}$, $oriV_{E.c.}$<br>pJC1-*nupG* LF-BcuI-*nupG* RF_1S derivative with an inserted *neoR*-oopT and an restored BcuI site.<br>Helper plasmid for reporter integration in *E. coli* | This work |
| pJC1-nupG LF-*neoR*-oopT-$P_{J23150}$-*gfpmut1*-*nupG* RF_3S | $Kan^R$, $oriV_{C.g.}$, $oriV_{E.c.}$<br>pJC1 derivative carrying the integration cassette for insertion of the *neoR* and $P_{J23150}$-*gfpmut1* sequence, separated by the oopT terminator.<br>$P_{J23150}$-*gfpmut1*-specific plasmid for integration in *E. coli*. | This work |
| pK18-*mobsacB*-cg1121/22-Lrp-sensor | $Kan^R$, $oriV_{E.c.}$, *sacB*, *lacZα*<br>Plasmid for genomic integration of the Lrp-sensor in the intergenic region of cg1121-cg1122 in *C. glutamicum*. | (Mustafi, Grünberger et al. 2014) |
| pK19-*mobsacB* | $Kan^R$, $oriV_{E.c.}$, *sacB*, *lacZα*<br>Plasmid for allelic exchange in *C. glutamicum* | (Schäfer, Tauch et al. 1994) |
| pSM24 | $kan^R$, pCLTON1$^{TS}$ (RepA$^{P47S}$) (#d) derivative containing the *dcas9* gene | Gift from S. Matamouros |
| pK19-*mobsacB*-cg1121/22_$P_{tet}$-*e2-crimson* | $Kan^R$, $oriV_{E.c.}$, *sacB*, *lacZα*<br>pK19-*mobsacB* derivative for inserting the e2-crimson gene under control of the constitutive $P_{tet}$ promoter (-*tetR*, constitutive) in the cg1121/22 intergenic region | This work |
| pPREx2-MDtxR-GA-G149 | $Kan^r$, pPREx2 derivative carrying the matryoshka cassette, encoding for the cpsfGFP and nested LSSmApple inserted into the respective insertion sites of DtxR (I138-V150) under the control of $P_{tac}$ promoter | (Papadopoulos, Anlauf et al. 2026) |
| pK19*mobsacB-strep-mcherry-dps* | $Kan^r$, pK19*mobsacB* derivative derivative encoding for the fusion protein mCherry-Dps | Sundermeyer, unpublished |
| pET24b-NHis | $Kan^r$, | Hentschel, unpublished |
| pPREx2-10xHis-TEV-MDtxR-GC-G149 | $Kan^r$, pPREx2 derivative carrying the matryoshka cassette, encoding for the cpsfGFP and nested mCherry inserted into the | This work |

| | | |
|---|---|---|
| | respective insertion sites of DtxR (I138-V150) with an N-terminal 10xHis tag and TEV cleavage site under the control of $P_{tac}$ promoter | |
| pRK2013 | *Kan^r^*, *oriV*(RK2/ColE1) -mob+ tra+ | Figurski and Helinski (1979) |
| pTNS1 | *Amp$^R$*, *oriV*(R6K), *TnSABC+D* operon | Choi, Gaynor et al. (2005) |
| mTn7_mCherry | *Kan$^r$*, Gm$^r$, *oriV* (R6K), pBG-derived, promoter Pem7, *mCherry* | Paik et al. unpublsihed |
| mTn7_mTagBFP | *Kan$^r$*, Gm$^r$, *oriV* (R6K), pBG-derived, promoter Pem7, *mTagBFP* | This study |
| mtn7_Ptac_IronSensR | *Kan$^r$*, Gm$^r$, *oriV* (R6K)*, Ptac, lacI, IronSensR* | Papadopoulos, Anlauf et al. (2026) |
| mtn7_Ptac_IronSensR-mCherry | *Kan$^r$*, Gm$^r$, *oriV* (R6K), *Ptac, lacI, IronSensR-mCherry* | This study |

**Table S5: Primers used in this study.**

| # | Sequence (5´-3´) | Purpose |
|---|---|---|
| **Primers for plasmid construction** | | |
| 1145 | AGCGACGCCGCAGGGGGATCCGGCGCAGGACATCACATTGCG | Construction of pJC1-Δ*entC*_disruption-cassette |
| 1146 | CGAAGCAGCTCCAGCCTACAATCATCCTCCACAAAATGATAAAGGC | |
| 1147 | TGTAGGCTGGAGCTGCTTCG | Construction of pJC1-Δ*entC*_disruption-cassette, sequencing of the genomic *entC* deletion |
| 1148 | CGGGGATCCGTCGACCTG | Construction of pJC1-Δ*entC*_disruption-cassette, sequencing of final plasmid |
| 1149 | CAGGTCGACGGATCCCCGCATGTTGAACGTTTTTGGATTGCATTAA | Construction of pJC1-Δ*entC*_disruption-cassette |
| 1150 | AAAACGACGGCCAGTACTAGCTACACGCGAGGTTATCCGC | |
| 1112 | AGCGACGCCGCAGGGGGATCCGATGTTCCTGATGATGACTAACGG | Construction of pJC1-*nupG* LF-BcuI-*nupG* RF_1S |
| 1114 | CGCAAAGAAAAACGGGTCGCC | |
| 1115 | AAAACGACGGCCAGTACTAGGGCGTGCTTTGCGTGGTACC | |
| 1604 | GGCGACCCGTTTTTCTTTGCGACTAGTTAATTAGTGGCTAACCGTCTG TGTG | |
| 1633 | GTGCACAAGGGATGTTCCTGATG | |
| 1634 | GTTGGTGCGGATTCGTGATGC | |
| 1605 | GACGGTTAGCCACTAATTAACTAGGTGTAGGCTGGAGCTGCTTC | Construction of pJC1-*nupG* LF-*neoR*-oopT-BcuI-*nupG* RF_2S |
| 1615 | GACCCGTTTTTCTTTGCGACTAGACTAGTAAAAAACGCCCGGCGGCA ACCGAGCGTTCTGAATTACTTAGTCTAGCTTACCTTAGTTCCTATTCC GGATCCGTCGACCTGCAGTT | |
| 1629 | CCCGTTTTTCTTTGCGACTAGACTAGCTATTTTTTGTATGGTTCATCCA TGCCATG | Construction of pJC1-nupG LF-*neoR*-oopT-$P_{J23150}$-*gfpmut1*-*nupG* RF_3S |
| 1630 | CGGTTGCCGCCGGGCGTTTTTTACTAGTTTACGGCTAGCTCAGTCCTA GGTATTATACTAGTTGAACTTTAAGAAGGAGATATCATATGGGTACC CTGCAGATGAGTAAAG | |
| 1592 | CAAACTGTCATTGCCATACGCGAATTCGTTGAACTAATGGGTGCTTTA GTTGAAG | Construction of pK19-*mobsacB*-cg1121/22_$P_{tet}$-*e2-crimson* |
| 1593 | TGATATCTCCTTCTTAAAGTTCATGCAGGTGTATCAACAAGCTGGG | |
| 259 | TGAACTTTAAGAAGGAGATATCATATGGATAGCACTGAGAACGTCAT C | |
| 1595 | CTATTTAAGAAGTTTAAATTGTGTCCATGAGTTCGCTCGACTACTGGA ACAGGTGGTGG | |
| 264 | cacaagcttaattaaGgaattcgagctcgg | Constuction of mTn7_mTagBFP |
| 265 | aatcagctcgctcattagaaaacctccttagcatg | |
| 266 | taaggaggttttctaatgagcgagctgattaagg | |
| 267 | ccgagctcgaattcCttaattaagcttgtgcccc | |
| 292 | CTGTACGGCGGCACCatggtgagcaagggc | Construction of mtn7_Ptac_IronSensR-mCherry |
| 293 | GCTGGCGCTGCCGCCcttgtacagctcgtccat | |
| 294 | gacgagctgtacaagGGCGGCAGCGCCAGC | |
| 295 | gcccttgctcaccatGGTGCCGCCGTACAG | |
| **Construction of pPREx2-MDtxR-GC-G149 (IronSenseR-II)** | | |
| Matr-S208 fw | CAAGTTGGACATCACCTCCCACAACGAGGACTAC | Construction of IronSenseR-mCherry |
| pPREx2 rv | ATGTATATCTCCTTCTGCAGGC | |
| 10xHisTEV fw | ATGCCTGCAGAAGGAGATATACATATGAGCAGCCATCATCATC | |
| TEV rv | CGGTGGTATCGACCAGATCCTTCATGCTAGCGCCCTGAAAATAC | |
| Matr-adj fw | ATGAAGGATCTGGTCGATAC | |

| | | |
|---|---|---|
| Matr rv | ATCCTCCTCGCCCTTGCTCACGGTGCCGCCATACAGTTC | |
| mCherry fw | ACTGTATGGCGGCACCGTGAGCAAGGGCGAGGAG | |
| mCherry-short rv | AGTCCTCGTTGTGGGAGGTGATGTCCAACTTGATG | |
| **Confirmation of integration by colony-PCR** | | |
| Tn7R | CACAGCATAACTGGACTGATTTC | Choi and Schweizer (2006) |
| PglmS-down | GCACATCGGCGACGTGCTCTC | |
| Seq1-MDtxR-GC-G149 fw | GCCGACATCATAACGGTTCTGG | Sequencing primers for IronSenseR-mCherry |
| Seq2-MDtxR-GC-G149 rv | CTTGTACAGCTCGTCCATGCCGC | |
| Seq3-MDtxR-GC-G149 rv | AGACCGCTTCTGCGTTCTG | |
| **Universal primers for plasmid sequencing** | | |
| R12 | CAGGGACAAGCCACCCGCACA | Universal sequencing primers of pJC1-based plasmids |
| R13 | GGAAGCTAGAGTAAGTAGTTCGC | |
| 920 | CATCGGCTCGTATAATGTGTGG | Universal sequencing primers of pEKEx2-based plasmids |
| 891 | GAAAATCTTCTCTCATCCGCC | |
| M19 | CGCCAGGGTTTTCCCAGTCAC | Universal sequencing primers of pK19-*mobsacB*-based plasmids |
| M20 | AGCGGATAACAATTTCACACAGGA | |
| **Primers for sequencing of genomic insertions/deletions** | | |
| 89 | CGCGCGATAAATGTGCGTTTGAG | Sequencing of the genomic insertions at integration locus cg1121/22 |
| 90 | CAAGGCCTAGTGTTCACTCAGG | |
| 921 | GTACCAGCGCGGTTTCACCAG | Sequencing of the genomic *entC* deletion |
| 922 | CGACATGCTGCGAGTGACAGC | |
| 633 | CAGGTCGACTCTAGAGGATCCAGAAGTTCGCTCAGCACCTTC | Sequencing of the $P_{J23150}$-*gfpmut1* insertion downstream of *nupG* |
| 1148 | CGGGGATCCGTCGACCTG | |
| 1629 | CCCGTTTTTCTTTGCGACTAGACTAGCTATTTTTTGTATGGTTCATCCATGCCATG | |
| 1634 | GTTGGTGCGGATTCGTGATGC | |

**Table S6: Description of all set parameters and initial concentrations used for the modelling of microfluidic co-culture growth.**

| Parameter | Description | Value | Unit | Source |
|---|---|---|---|---|
| $\mu_{max-E.coli}$ | Maximum growth rate of *E. coli* Wildtype in monoculture | 0.25 | 1/h | This work |
| $\mu_{min-E.coli}$ | Maximum growth rate of *E. coli* Δ*entC* in monoculture | 0.15 | 1/h | This work |
| $\mu_{max-C.glutamicum}$ | Maximum growth rate of *C. glutamicum* on glucose | 0.4 | 1/h | (Unthan, Grünberger et al. 2014) |
| $k_{S,X2}$ | kS value of enterobactin uptake by *E. coli* | 0.2678 | mg/L | Thulasiraman, Newton Salete et al. (1998) |
| $k_{S,X1}$ | kS value of enterobactin uptake by *C. glutamicum* | 10 x $k_{S-E-E}$ | mg/L | Adjusted to check our hypothesis |
| $q_{max_{p-1-E.coli}}$ | Production of abstract enterobactin-iron product at maximum growth rate of *E. coli* | 90% of $k_{S-E-C}$ | mg/L*h | Adjusted to check our hypothesis |
| $q_{max_{c-1-E.coli}}$ | Consumption of abstract enterobactin-iron product at maximum growth rate of *E. coli* | 0.4 x of $q_{max_{p-1-E.coli}}$ | mg/L*h | Adjusted to check our hypothesis |
| $q_{max_{c-1-C.glut}}$ | Consumption of abstract enterobactin-iron product at maximum growth rate of *C. glutamicum* | 1.6 x of $q_{max_{c-1-E.coli}}$ | mg/L*h | Adjusted to be the same differences as the growth rates of the organisms |
| $a_{\emptyset}$ | Average cell area of a single cell | 2.0 | $\mu m^2$ | This work |
| $c_{cdw-c}$ | Average mass of cell dry weight by volume unit of cell | 250 | g/L | Smaluch, Wollenhaupt et al. (2023) |
| $V_{M-P-C}$ | Volume of a microfluidic perfusion chamber | 6000 | $\mu m^3$ | This work |
| $d$ | Cell diameter and chamber height | 1 | µm | This work |
| $P_0$ | Enterobactin concentration at the start | 15 | mg/L | Adjusted to check our hypothesis |
| $X_{Total-t_0}$ | Total cell dry weight concentration at the start of the simulation | Varied | g/L | Varied to account for different start cell areas and compositions |

**Table S7: Nomenclature of modelled parameters.**

| Parameter | Descripton | Unit |
|---|---|---|
| $\mu_{max-E.coli}$ | Maximum growth rate of *E. coli* Wildtype in monoculture | [1/h] |
| $\mu_{min-E.coli}$ | Maximum growth rate of *E. coli* Δ*entC* in monoculture | [1/h] |
| $A_{cell}$ | Average area of a single cell | [µm$^2$] |
| $A_{cylinder}$ | Projected area of the cylindrical part of the average cell | [µm$^2$] |
| $A_{sphere}$ | Projected area of the spherical part of the average cell | [µm$^2$] |
| $A_t$ | Observed total projected cell area at timepoint *t* for an entire chamber | [µm$^2$] |
| $c_{A\rightarrow V}$ | Factor to calculate cell area to cell volume. Describes how much cell volume is existing per µm$^2$ of projected cell area. | [µm$^3$/µm$^2$] |
| $c_{cdw-c}$ | Average mass of cell dry weight by volume unit of cell | [g/L] |
| $d$ | Cell diameter and chamber height | [µm] |
| $k_{S-E-C}$ | kS value of enterobactin-iron product uptake by *C. glutamicum* | [mg/L] |
| $k_{S-E-E}$ | kS value of enterobactin-iron product uptake by *E. coli* | [mg/L] |
| $L_{cylinder}$ | Length of the cylindrical part of the average cell | [µm] |
| $P_{Enterobactin}$ | Concentration of abstract enterobactin-iron product | [g/L] |
| $P_0$ | Enterobactin-iron product concentration at the start | [mg/L] |
| $q_{max_{c-1-C.glut}}$ | Consumption of abstract enterobactin-iron product at maximal growth rate of *C. glutamicum* | [mg/(L*h)] |
| $q_{max_{c-1-E.coli}}$ | Consumption of abstract enterobactin-iron product at maximal growth rate of *E. coli* | [mg/(L*h)] |
| $q_{max_{p-1-E.coli}}$ | Production of abstract enterobactin-iron product at maximal growth rate of *E. coli* | [mg/(L*h)] |
| $V_{sphere}$ | Volume of the spherical part of the cell | [µm$^3$] |
| $X_{C.glutamicum}$ | Cell dry weight concentration of *C. glutamicum* | [g/L] |
| $X_{E.coli}$ | Cell dry weight concentration of *E. coli* | [g/L] |
| $X_{Total}$ | Total cell dry weight concentration, combination of both organisms | [g/L] |
| $X_{Total-t_0}$ | Total cell dry weight concentration at the start of the simulation | [g/L] |
| $X_t$ | Cell dry weight concentration at time point t | [g/L] |

**Formula S1: Derivation of conversion factor between measured cell area and volume of cells in the microfluidic system.**

Three conversion steps were performed to convert observed cell areas into simulated cell dry weight concentrations. First the volume of all cells in the system is calculated by the measured cell area. Cells are assumed to be of the same average cell area and of the same geometry, which does not change over time. The cell morphology was described by a cylinder capped by two hemispheres. The diameter of the cell is assumed to be same as the height of the microfluidic system. A conversion factor is then used to calculate any measured projected cell area to cell volume.

$$A_{sphere} = \pi \times \frac{d}{2}^2 \tag{1}$$

$$A_{cylinder} = A_{cell} - A_{sphere} \tag{2}$$

$$L_{cylinder} = \frac{A_{cylinder}}{d} \tag{3}$$

$$V_{sphere} = \frac{4}{3} \times \pi \times \frac{d}{2}^3 \tag{4}$$

$$V_{cylinder} = \frac{d}{2}^2 \times \pi \times L_{cylinder} \tag{5}$$

$$V_{cell} = V_{sphere} + V_{cylinder} \tag{6}$$

$$c_{A \to V} = \frac{V_{cell}}{A_{cell}} \tag{7}$$

$$c_{A \to V} = \frac{\frac{4}{3} \times \pi \times \frac{d}{2}^3 + \frac{d}{2}^2 \times \pi \times \frac{A_{cell} - \pi \times \frac{d}{2}^2}{d}}{A_{cell}} \tag{8}$$

The calculated cell volume is then processed with an average value for cell dry weight per volume to cell dry weight. Lastly the cell dry weight concentration in a chamber is calculated by using the volume of a microfluidic chamber.

$$X_t = \frac{A_t \times c_{A \to V} \times c_{cdw-c}}{V_{M-P-C}} \tag{9}$$

$$X_{total} = X_{C.glut} + X_{E.coli} \tag{10}$$
